# The RNA-binding protein TRIM71 is essential for hearing in humans and mice and regulates the timing of auditory sensory organ development

**DOI:** 10.1101/2025.05.29.656801

**Authors:** Xiao-Jun Li, Lin Li, Charles Morgan, Phan Q. Duy, Lale Evsen, Le T. Hao, Roxane Machavoine, Kahina Belhous, Sylvain Ernest, Françoise Denoyelle, Cyril Mignot, Frederic Brioude, Marine Parodi, He Huang, Prathamesh T Nadar Ponniah, Waldemar Kolanus, Kristopher T. Kahle, Sandrine Marlin, Angelika Doetzlhofer

## Abstract

The RNA-binding protein TRIM71 is essential for brain development, and recent genetic studies in humans have identified *TRIM71* as a risk gene for congenital hydrocephaly (CH). Here, we show that mono-allelic missense mutations in *TRIM71* are associated with hearing loss (HL) and inner ear aplasia in humans. Utilizing conditional *Trim71* knockout mice carrying a CH and HL-associated mutation, we demonstrate that loss of TRIM71 function during early otic development (embryonic day 9-10) causes severe hearing loss. While inner ear morphogenesis occurs normally in *Trim71* knockout mice, we find that early otic loss of TRIM71 function disrupts the highly stereotyped timing of cell cycle exit and differentiation within the inner ear auditory sensory organ (cochlea), resulting in the premature formation and innervation of mechano-sensory hair cells. Transcriptomic profiling of *Trim71* deficient cochlear progenitor cells identifies *Inhba* and *Tgfbr2* as targets of TRIM71 repression, and our analysis of *InhbaTgfbr1* double knockout mice indicates that TRIM71 maintains hair cell progenitors in a proliferative and undifferentiated state by restricting TGF-β-type signaling. Characterization of hair cells and their associated neurons in adult *Trim71* knockout mice revealed abnormally short inner hair cell stereocilia, reduced pre-synaptic terminals, and neuronal degeneration in the outer hair cell region, providing a basis for the observed hearing deficits in *Trim71* knockout mice.

## INTRODUCTION

The inner ear cochlea contains a highly specialized sensory organ dedicated to detecting sound. The core functional unit of this sensory organ comprises mechanoreceptors-termed hair cells, which are positioned atop supporting cells and are innervated by afferent spiral ganglion neurons (SGNs). Hair cells, supporting cells, and SGNs are generated in a highly stereotyped manner that requires strict spatial and temporal control of cell proliferation and differentiation (reviewed in ^1^). Recent functional studies have revealed that the RNA-binding protein LIN28B and members of the *let-7 (lethal)* family of miRNAs play a central role in timing cell cycle exit and differentiation of hair cell and supporting cell progenitors (termed pro-sensory cells) in the murine ^2^ and avian auditory sensory organ ^3^. The *let-7* miRNAs and their mutual antagonist LIN28B are part of an evolutionary highly conserved network of genes, first identified in *C. elegans*, which controls developmental timing and stemness ^4,5^. While LIN28B promotes stemness, *let-7* miRNAs promote cell cycle withdrawal and differentiation through targeting growth-related genes, including *Lin28b* itself ^6^. Another well-known *let-7* target and evolutionarily conserved gene that promotes stemness is *Trim71* (also referred to as *Lin41*) ^5,7^. The expression of *Trim71* peaks during early embryonic development ^8,9^, and *Trim71* continues to be expressed in various stem cell populations, such as germ cells, throughout later developmental stages^10^. Highlighting TRIM71’s importance for stemness, TRIM71 has been found to enhance the reprogramming of human fibroblasts into induced pluripotent stem cells (iPSCs) when co-expressed along with pluripotency factors OCT4, SOX2, and KLF4 ^11^. TRIM71 belongs to the tripartite-motif (TRIM)-NHL protein family, characterized by an N-terminal RING domain with intrinsic ubiquitin E3 ligase activity and a C-terminal NHL domain responsible for RNA-binding reviewed in ^12^. Genetic and biochemical studies showed that TRIM71 cooperates with core elements of the miRNA-induced silencing complex to repress the translation of targeted mRNAs and facilitate their degradation ^13–16^. During early embryonic development, TRIM71 is instrumental in preventing the premature activation of neurogenic genes, and *Trim71* gene trap ^8,9^ and knockout mice ^14^ have severe neural tube closure defects due to decreased cell proliferation and premature onset of neural differentiation. In humans, miss-sense mutations in the *TRIM71* gene lead to a neurodevelopmental syndrome characterized by ventriculomegaly and hydrocephalus ^17–20^. Loss of function studies in mice revealed that brain abnormalities arise from the premature differentiation of neuroepithelial cells, adversely affecting cortical neurogenesis and subsequent cerebrospinal fluid biomechanics ^19^. We now show that CH-causing miss-sense mutations in *TRIM71* are linked to hearing loss (HL) in humans. Using conditional *Trim71* knockout mice that carry the mouse equivalent of the human CH and HL-associated mutation, we demonstrate that early otic loss of TRIM71 function results in severe hearing loss. Transcriptomic and functional data demonstrate that TRIM71 times cell cycle exit and differentiation within the auditory sensory epithelium by repressing *Inhba* and *Tgfbr2* expression, two key components of TGF-β-type signaling. Characterization of hair cells and their associated neurons in adult Trim71 knockout mice revealed abnormally short hair cell stereocilia, reduced pre-synaptic terminals, and neuronal degeneration in the outer hair cell region. In sum, our research identifies TRIM71 as a syndromic HL gene and critical regulator of auditory-sensory development.

## RESULTS

### Mono-allelic missense mutations in *TRIM71* are associated with hearing loss in humans

Recent human genetics studies identified *TRIM71* as a novel disease gene in congenital hydrocephaly ^17–20^. A subset of patients with *de novo* or transmitted mono-allelic *TRIM71* mutations reported hearing problems and underwent further testing (Table 1). These patients have been reported in previous studies ^19,20^. Three out of nine patients reported a hearing loss phenotype, and for two of these patients, details regarding clinical workup of hearing loss were available (Patients KCHYD673-1 and 18CY000656, also referred to as “patient 1” and “patient 2”, respectively, in the rest of the manuscript). Patient 1, a female who harbors a p.(Gln334Arg) mutation in TRIM71, which impairs the native subcellular localization of TRIM71 to P-bodies ^20^, presented with a congenital hypothyroidism with ectopic thyroid, a right mild sensorineural hearing loss, bilateral chorioretinal coloboma, bilateral renal cysts, axial hypotonia, brachydactyly clinodactyly of the 5th fingers, a square face, round external ears without lobule and bifid uvula at 7 months of age. At 5 years old, the hearing loss has progressed to a mixed severe right hearing loss (Figure S1A). Brain and internal auditory canals MRI showed bilateral lateral semicircular canals malformation with normal cochlea and olfactory bulbs (Figure S2A-C). She had motor delay with abnormal vestibular tests without other neurodevelopmental impairment. Patient 2, a female who harbors a missense mutation in TRIM71’s NHL domain [p.(Arg608His)], which impairs RNA binding ^19^, presented with a mixed severe left and a moderate conductive right hearing loss at 6 years old (Figure S1B). Temporal bone CT scans showed unilateral left semicircular canal malformation with normal cochlea, bilateral external ear canal hypoplasia, and ossicular malformations (Figure S2D-K). She also presented with speech and language delay, hydrocephalus, brachymesophalangia of the 5th left finger, ventricular septal defect, and only 4 lumbar vertebrae. These clinical findings suggest a potential link between *TRIM71* dysfunction and human hearing loss.

**Table 1.**
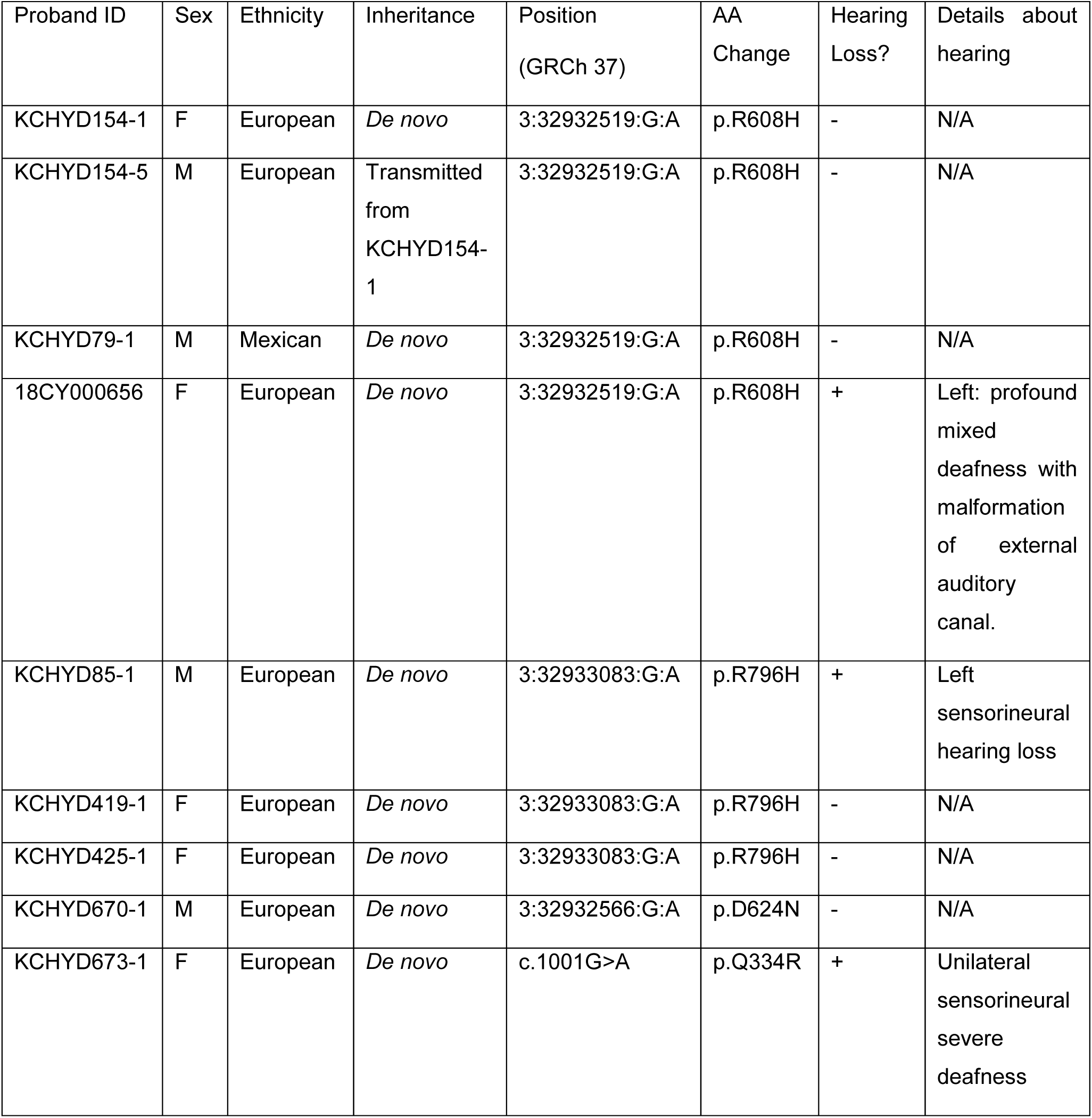
Genetic and clinical phenotypes of patients with mutations in *TRIM71*.

### *Trim71* is expressed in otic neural-sensory progenitor cells

To gain insights into how miss-sense mutations in *Trim71* may cause conductive and or sensorineural hearing loss, we utilized published single-cell RNA sequencing data to examine *Trim71* expression in the developing mouse inner ear at stages E9.5, E11.5, and E13.5 ^21^ using Gene Expression Analysis Resource (gEAR) portal ^22^. The inner ear epithelium and its innervating afferent neurons derive from the otic placode located adjacent to the hindbrain. By E9.5, the otic placode-derived cells have formed an otocyst and by E11.5, many of the neuroblasts have delaminated and coalesced to form the otic ganglion, which later subdivides into cochlear (spiral ganglion) and vestibular ganglion, and by E13.5, both vestibular and auditory sensory epithelia are specified (reviewed in ^23^).

We found that at stage E9.5, *Trim71*, similar to *Lin28b*, was broadly expressed in both otic and peri-otic cells, with high expression in otic epithelial cells marked by *Tbx2* and *Lfng* expression (Figure 1A). At E11.5 and E13.5, *Trim71* expression became more refined, co-expressing with *Lin28b* in otic epithelial and ganglion cells at E11.5, and at E13.5 in inner ear sensory and cochleovestibular ganglion cells (Figure 1A). We independently confirm *Trim71’*s early otic expression in stage E9.5 mouse tissue using anti-TRIM71 immuno-staining (Figure 1B) and RNA in situ hybridization (ISH) (Figure 1C). *Trim71* transcript was highly expressed in the developing otocysts, particularly in the neural sensory competence domain marked by *Lfng* expression ^24^ and, consistent with previous reports, in the ventricular zone of the adjacent hindbrain ^19^ (Figure 1C). Immunostaining with anti-TRIM71 antibody showed broad expression of TRIM71 protein in otic and peri-otic tissue and confirmed previous findings that TRIM71 protein resides in the cytoplasm (Figure 1B). Next, we used quantitative PCR (qPCR) to analyze changes in *Trim71* expression prior (E13.5) and during cochlear hair cell differentiation (E14.5-E16.5). As a control, we also examined the expression of pro-sensory genes *Isl1* ^24^, *Hmga2,* and *Lin28b* ^2^, along with the hair cell marker *Atoh1*. The transcription factor ATOH1 is highly expressed in nascent hair cells, and induction of *Atoh1* expression within the cochlear sensory epithelium marks the onset of hair cell differentiation and cochlear differentiation in general ^25^. Like *Isl1*, *Lin28b,* and *Hmga2*, the expression of *Trim71* was highest prior to differentiation (E13.5). However, while *Isl1,* which continues to be expressed in supporting cells, only modestly declined during differentiation, the expression of *Trim71*, *Lin28b,* and *Hmga2* sharply declined at the onset of differentiation (E13.5-E14.5), and *Trim71* was near undetectable in differentiating cochlear epithelia (E15.5-E16.5) (Figure 1D).

**Figure 1.**
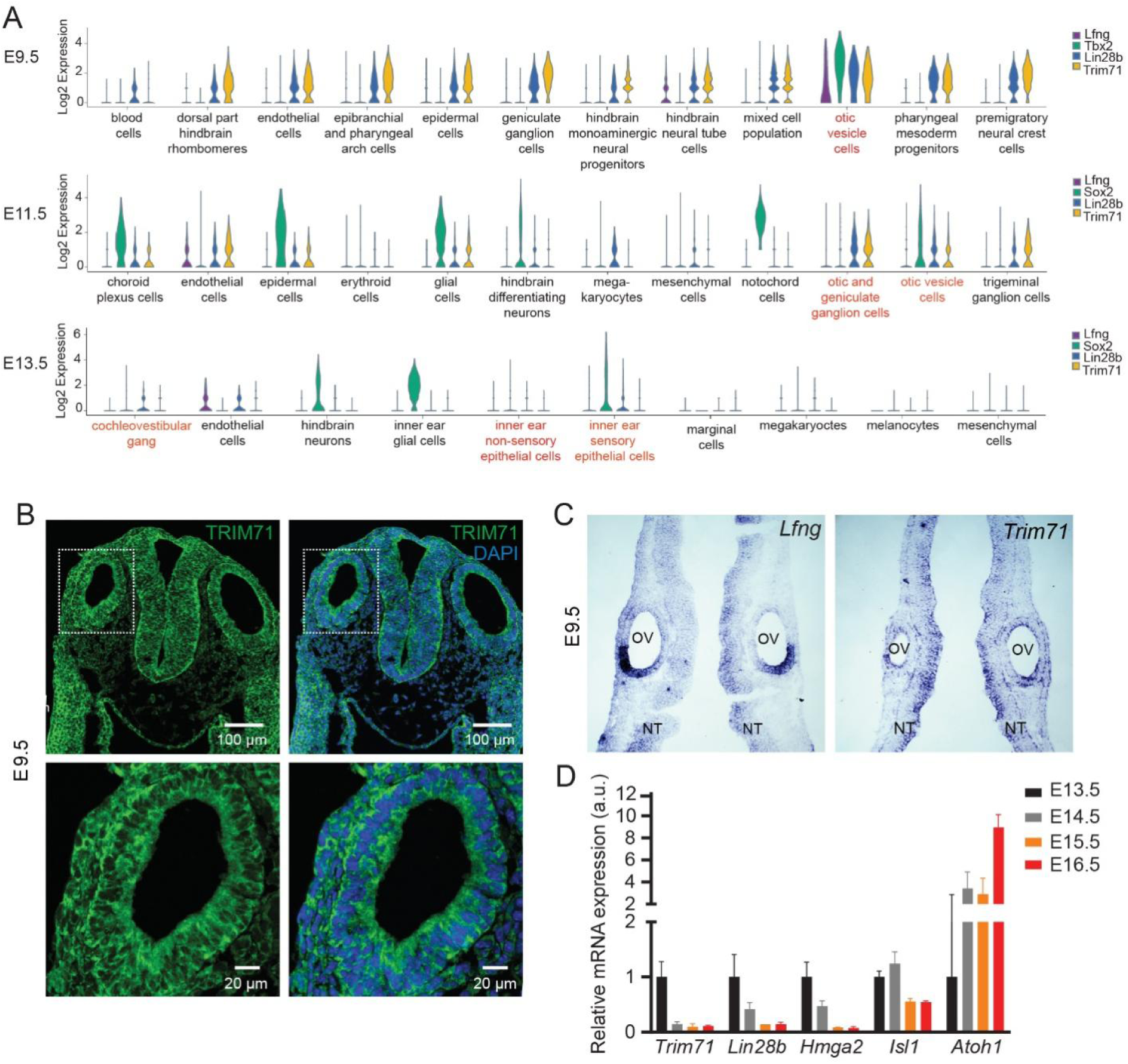
*Trim71* is expressed in otic and neuro-sensory progenitor cells. (A) *Trim71* and *Lin28b* mRNA expression in otic and periotic cells stages E9.5, E11.5, and E13.5. Violin plots were generated from previously published scRNA-sequencing data sets. *Tbx2*, *Lfng,* and *Sox2* served as marker genes. (B) Otic and peri-otic TRIM71 protein expression in stage E9.5 mouse embryo. (C) Otic and peri-otic *Trim71* and *Lfng* mRNA expression in stage E9.5 mouse embryo. (D) RT-qPCR of *Trim71, Lin28b, Hmga2, Isl1, and Atoh1* in cochlear epithelia isolated from stage E13.5, E14.5, E15.5, and E16.5 mouse embryos. Bar graph showing mean ± SD, n=3 biological replicates per group.

### *Trim71* knockout during early otic development impairs hearing in mice

We established *Trim71* mutant mouse models to examine the role of TRIM71 in inner ear development and hearing. Non-conditional *Trim71* homozygous mutant mice have severe neural tube closure defects and die mid-gestation ^8,14,26^. To circumvent embryonic lethality, we employed a conditional knockout approach using *Trim71* floxed (*Trim71^f/f^*) mice. In these mice, exon 4, which codes for the NHL domain, is flanked by LoxP sites, which, upon Cre-mediated recombination, renders the *Trim71* gene non-functional ^14^ (Figure 2A). To achieve broad yet stage-specific *Trim71* deletion, we used a doxycycline (dox) inducible Cre strategy (*R26^rtTA*M2^* and *TetO-Cre*). Additionally, we generated *Trim71* mutant mice (*Trim7^R595H/Δ^*), where one floxed allele was replaced by an allele that carries the murine homolog (R595H) of the human CH-and HL-associated missense mutation in *TRIM71* (R608H) ^19^ (Figure 2B). We validated our loss-of-function strategy in pilot experiments where dox was administrated at E5.5 (Figure 2C).

**Figure 2.**
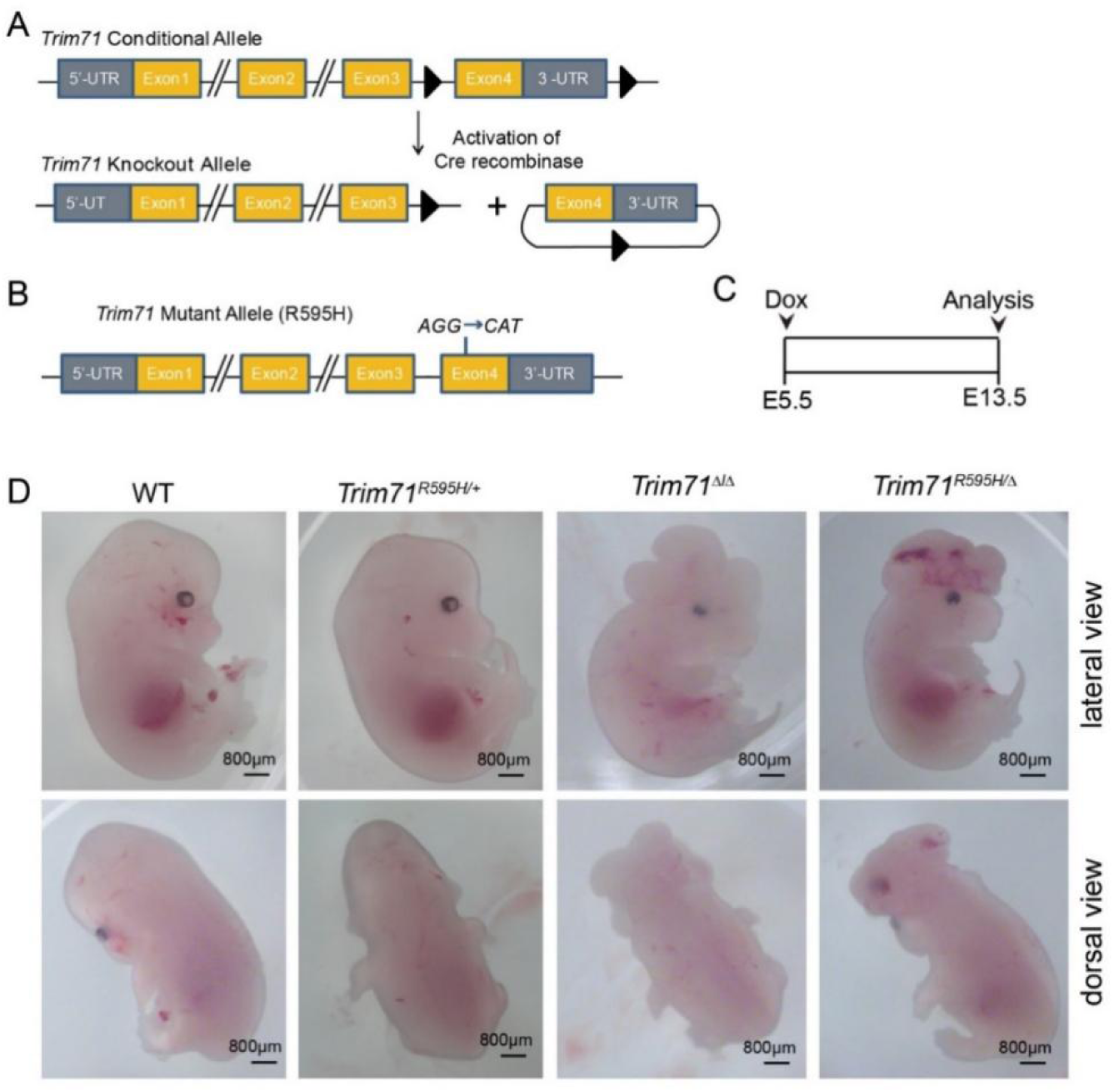
*Trim71* homozygous mutant mice exhibit neural tube closure defects. (A) Schematic of *Trim71* conditional and knockout allele. (B) Schematic of *Trim71* mutant allele. (C) Experimental design for D. Doxycycline-inducible Cre strategy (*R26^rtTA*M2^*; *TetO-Cre*) was used to knockout *Trim71* conditional allele(s) at E5.5. (D) *Trim71* knockout mice and *Trim71* mutant mice have severe neural tube closure defects. Shown are bright field images of stage E13.5 mice that are wild type for *Trim71* (WT), heterozygous for mutant allele (*Trim71^R595H/+^*), homozygous for knockout allele (*Trim71^Δ/Δ^*) or carry mutant and knockout alleles (*Trim71^R595H/Δ^)*.

As expected, dox administration at E5.5 (pre-gastrulation) resulted in severe neural tube closure defects in *Trim71* knockout (*Trim7^Δ/Δ^*) or *Trim71* mutant (*Trim7^R595H/Δ^*) embryos (Figure 2D). A small fraction (<10%) of *Trim71* heterozygous mutant mice (*Trim7^R595H/+^*) displayed neuro-developmental defects, as reported by ^19^. However, the majority of *Trim71* heterozygous mutant (*Trim7^R595H/+^*) embryos developed normally and were indistinguishable from their wild-type littermates (Figure 2D). Furthermore, dox administration at E8.5 or later allowed *Trim71* knockout and *Trim71* mutant mice to survive into adulthood (Figure 3A). At one month, *Trim71* knockout and *Trim71* mutant (*Trim7^Δ/Δ^*and *Trim7^R595H/Δ^*) mice weighed significantly less than *Trim71* heterozygous mutant (*Trim7^R595H/+^*) or control (*Trim71*^f/f^) mice (Figure 3B). Additionally, *Trim71* knockout and *Trim71* mutant mice were smaller overall and had notably shorter tails than *Trim71* heterozygous mutant or control mice (Figure 3A).

**Figure 3.**
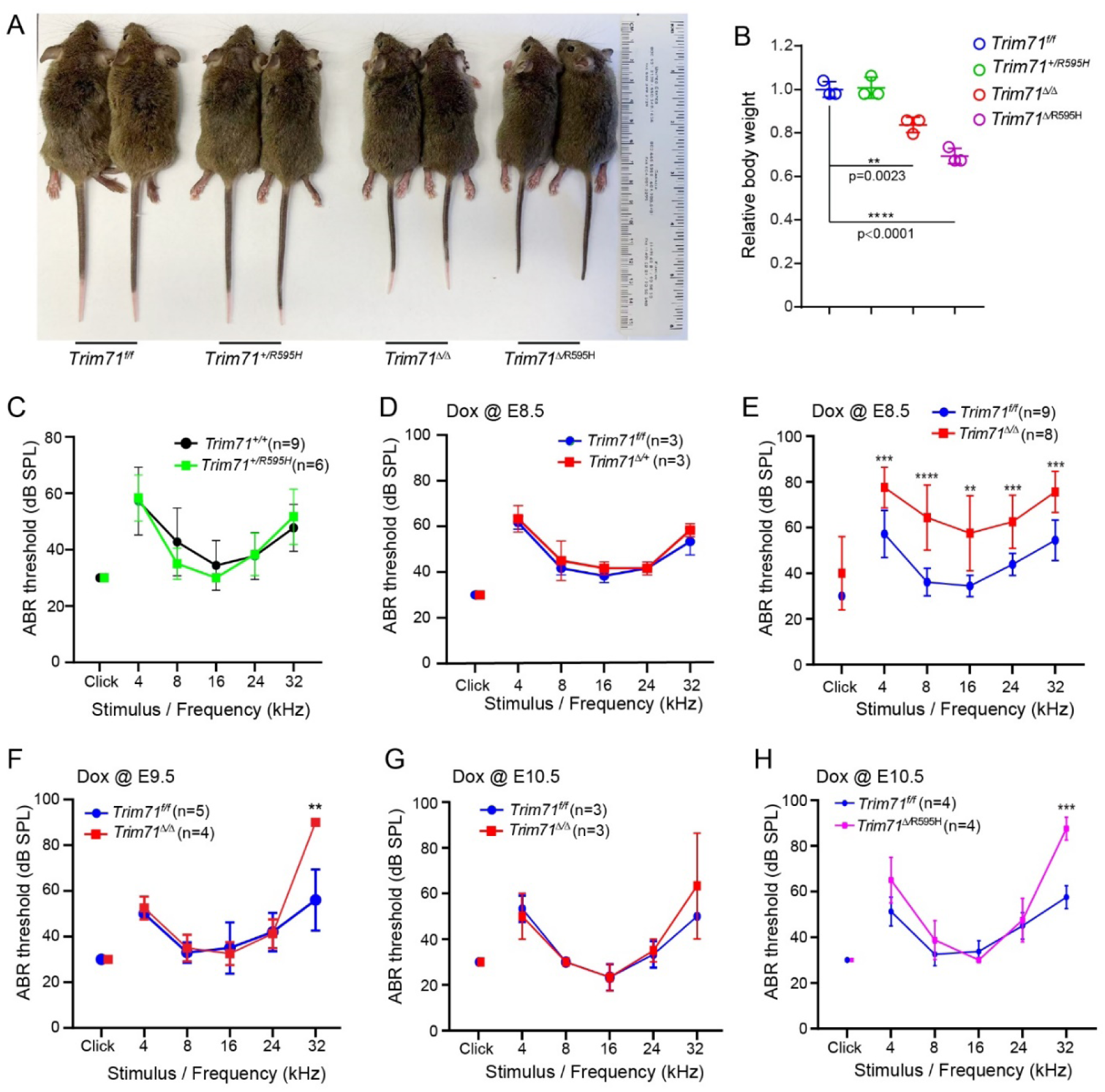
Early embryonic loss of *Trim71* impairs hearing in mice. Doxycycline (dox) inducible Cre strategy *(R26rtTA*M2; Tet-on-Cre*) was used in mice to conditionally delete *Trim71* during early otic development (E8.5-E10.5). (A) P30 *Trim71^Δ/Δ^* and *Trim71^R595H/Δ^* mice that received dox starting at E8.5 are reduced in size compared to *Trim71^f/f^*(WT, control) and *Trim71^R595H/+^* mice. (B) Quantification of the body weight in (A) (n=4 animals, two independent experiments). (C-H) Auditory Brainstem Response (ABR) thresholds were measured in 1-month old mice. (C) ABR thresholds were not elevated (normal hearing) in *Trim71^R595H/+^*mice compared to *Trim71^+/+^* wild type littermates. (D) ABR thresholds were not elevated in *Trim71^+/Δ^* mice compared to *Trim71^f/f^* (control) littermates. Dox was administered starting at E8.5 (n=3 animals per group). (E) ABR thresholds were elevated in *Trim71^Δ/Δ^* mice across all frequencies compared to *Trim71^f/f^* control littermates after dox was administered starting at E8.5 (n=9 animals in control group and n=8 animals in *Trim71^Δ/Δ^* group, three independent experiments). (F) ABR thresholds were elevated at high frequency (32 kHz) in *Trim71^Δ/Δ^*mice compared to control *Trim71^f/f^* littermates after dox was administered at E9.5 (n=5 animals in control group, n=4 animals in *Trim71^Δ/Δ^*group). (G) ABR thresholds were normal in *Trim71^Δ/Δ^* mice compared to *Trim71^f/f^* control littermates after dox was administered at E10.5 (n=3 animals in control group, n=3 animals in *Trim71^Δ/Δ^*group). (H) ABR threshold was significantly elevated at high frequency in *Trim71^Δ/R595H^* mice compared to *Trim71^f/f^* control littermates after the dox was administered at E10.5 (n=4 animals per group). One-way ANOVA with Tukey’s correction was used to calculate P values in B. Two-tailed, unpaired Student *t* test was used to calculate *P* values in C-H.\**P* < 0.05, \*\**P* < 0.01, \*\*\**P* < 0.001, and \*\*\*\**P* < 0.0001. All data are collected from at least two independent experiments.

Next, we used auditory brainstem response (ABR) threshold measures to characterize the hearing of *Trim71* homozygous and heterozygous knockout and mutant mice, as well as their control littermates (*Trim71*^f/f^ and *Trim71*^+/+^). Our analysis revealed that *Trim7^R595H/+^* mice had similar ABR thresholds to their wild-type littermates, indicating normal hearing (Figure 3C). Similarly, *Trim71* heterozygous knockout mice, with one floxed allele (*Trim71^+/Δ^*), exhibited normal hearing after receiving dox at E8.5 (Figure 3D). In contrast, *Trim71* homozygous knockout mice (*Trim7^Δ/Δ^*) had elevated ABR thresholds at low (4, 8 kHz), mid (16, 24 kHz), and high frequencies (32 kHz) compared to control littermates (*Trim71*^f/f^) after receiving dox at E8.5 (Figure 3E). Conditional deletion of *Trim71* after dox administration at E9.5 still impaired hearing at high frequency (Figure 3F, 32 kHz), but conditional deletion of *Trim71* after stage E10.5 had no adverse effect on hearing (Figure 3G). *Trim71* mutant mice that carried the R595 mutant allele and a floxed allele (*Trim7^△/R595H^*) displayed high-frequency hearing loss after receiving dox at E10.5, indicating that the R9595 miss-sense mutation impairs hearing (Figure 3H). In summary, our findings show that TRIM71 function during early otic development (E8.5-E10) is essential for proper hearing.

### *Trim71* knockout during early otic development results in premature pro-sensory cells cell cycle exit and differentiation

To gain insights into the genesis of the observed hearing deficits, we analyzed whether loss of TRIM71 during early otic development (dox starting between E8.5-E10.5) alters the behavior of auditory pro-sensory cells. Cochlear (auditory) hair cells and neighboring supporting cells derive from a pool of SOX2-expressing pro-sensory cells within the developing cochlear epithelial duct ^27,28^. Cochlear pro-sensory cells exit the cell cycle in an apical-to-basal gradient that in mice starts at around E12.5 and peaks at E13.5 ^29^. Following terminal mitosis, pro-sensory cells located at the cochlear mid-base start to differentiate at around E14.0, initiating a basal-to-apical wave of differentiation that reaches the cochlear apex around E18.5 ^25,30^. To capture the terminal mitosis of pro-sensory cells, we administered a single pulse of EdU at E12.5 (apex final mitosis) or E13.5 (mid-base final mitosis) and analyzed EdU incorporation in hair cells and supporting cells (Deiter’s cells and pillar cells) five days later.

We found that an EdU pulse at E12.5 labeled 2-fold fewer apical hair cells (Figure 4A, B) and apical supporting cells (Figure 4A, C) in E17.5 *Trim71^Δ/Δ^* mice than control littermates, indicating that in the absence of *Trim71* pro-sensory cells exit the cell cycle prematurely. Furthermore, we found that the length of the cochlear sensory epithelium in Trim71 knockout mice was about 20% shorter compared to control littermates (Figure 4D). A similar phenotype was observed with *Trim7^R595H/Δ^*mice. We found that an EdU pulse at E13.5 labeled 2-fold fewer hair cells in the cochlear base (Figure 4E, F) and more than 2-fold fewer supporting cells in the cochlear mid and apical region in *Trim7^R595H/Δ^* mice than control littermates (Figure 4E, G). However, loss of *Trim71* did not disrupt the apex-to-base gradient of pro-sensory cell cycle exit, and in both control and *Trim7^R595H/Δ^* mice, EdU incorporation in hair cells and supporting cells was highest in the cochlear base and lowest in the apex (Figure 4E-G). These results are consistent with our recent findings that cochlear epithelial cells isolated from stage E13.5 *Trim7 ^Δ/Δ^* or *Trim7^△/R595H^* mice (dox E5.5) proliferate at a significantly lower rate than cochlear epithelial cells isolated from control littermates when cultured as organoids *in vitro* ^31^.

**Figure 4.**
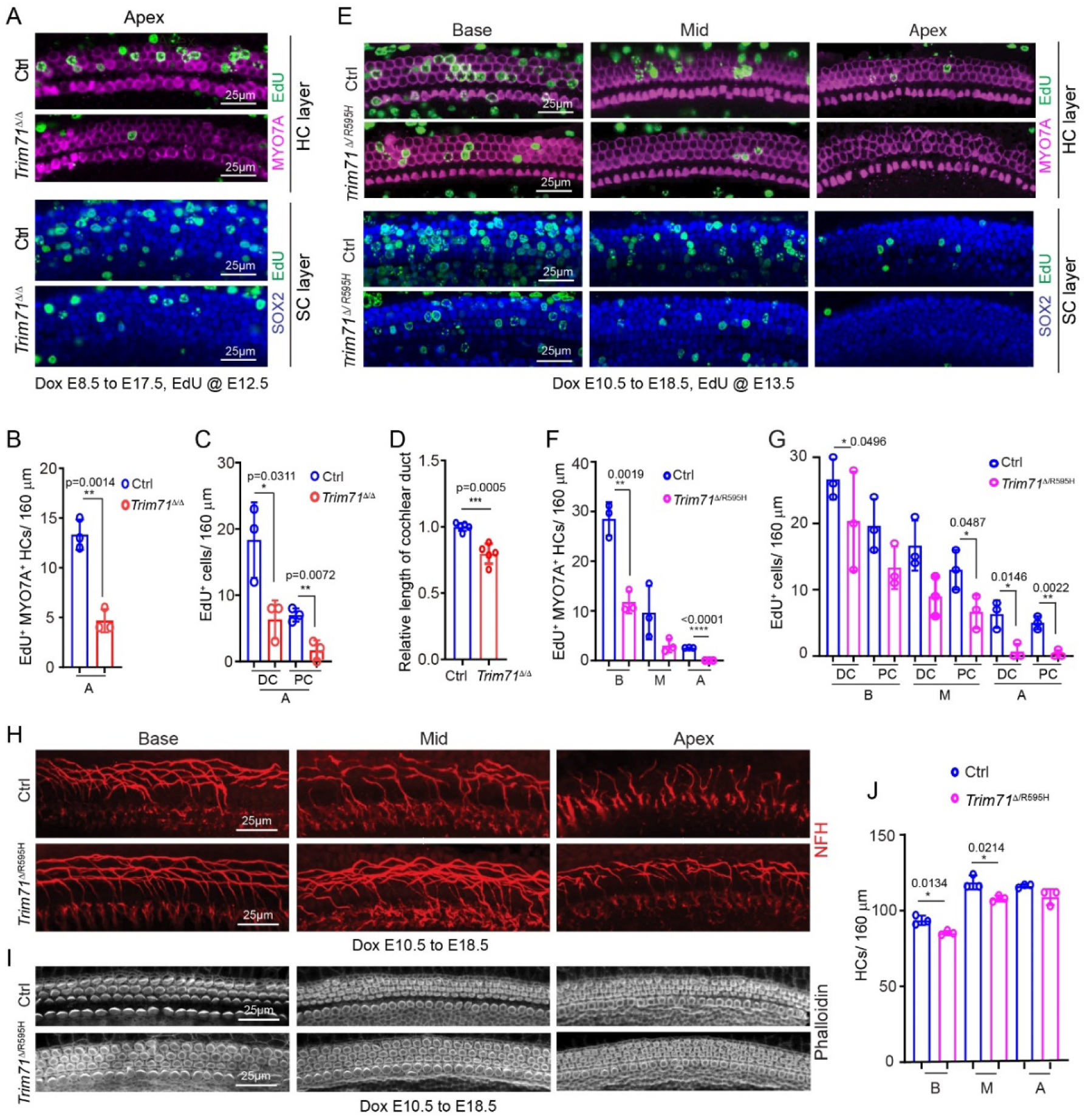
Loss of *Trim71* leads to premature cell cycle withdrawal and differentiation of auditory pro-sensory cells. (A-D) The timed-mated pregnant dam was fed doxycycline (dox)-containing feed starting at E8.5, a single EdU pulse was given at E12.5 and EdU (green) incorporation in *Trim71* knockout *(Trim71*^Δ*/*Δ^) mice and control *(Trim71^f/f^*) littermates was analyzed at E17.5. Cochlear length of *Trim71*^Δ*/*Δ^ mice and control *(Trim71^f/f^*) littermates was analyzed at P0. (A) Representative confocal images of EdU incorporation (red) in hair cells (MYO7A, magenta) and supporting cells (SOX2, blue) at the cochlear base, mid and apex in E17.5 *Trim71*^Δ*/*Δ^ mice and control littermates are shown. (B) Hair cell progenitors withdraw from the cell cycle prematurely in *Trim71*^Δ*/*Δ^ mice. Quantification of hair cell-specific EdU incorporation in (A) (n=3 animals per group, two independent experiments). (C) Supporting cell progenitors withdraw from the cell cycle prematurely in *Trim71*^Δ*/*Δ^ mice. Quantification of supporting cell-specific EdU incorporation in (A). Deiters cells (DC) and pillar cells (PC) are quantified (n=3 animals per group, two independent experiments). (D) Length of cochlear sensory epithelia in *Trim71* knockout (*Trim71*^Δ*/*Δ^) mice is shorter than in control mice (n=5 animals per group, two independent experiments). (E-G) The timed-mated pregnant dam was fed dox-containing feed starting at E10.5, EdU was injected at E13.5, and cochlear tissue of *Trim71^Δ/R595^* mice and *Trim71^f/f^* (control) littermates was collected and analyzed at E18.5. (E) Representative confocal images of EdU incorporation (red) in hair cells (MYO7A, magenta) and supporting cells (SOX2, blue) at the cochlear base, mid, and apex in E18.5 *Trim71^Δ/R595^*mice and control littermates are shown. (F) Hair cell progenitors withdraw from the cell cycle prematurely in *Trim71^Δ/R595^* mice compared to control littermates. Quantification of hair cell-specific EdU incorporation in (E) (n=3 animals per group, three independent experiments). (G) Supporting cell progenitors prematurely withdraw from the cell cycle in *Trim71^Δ/R595^* mice. Quantification of supporting cell-specific EdU incorporation in (E) (n = =3 animals per group, three independent experiments). (H) Outer hair cell innervation occurs prematurely in *Trim71^Δ/R595^* mice. Representative confocal images of neuronal innervation pattern (NFH, red) at the cochlear base, mid and apex of *Trim71^Δ/R595^* mice and control littermates are shown. (I) Hair cell stereocilia (phalloidin, grey) form prematurely in *Trim71*^Δ/R595^ mice. Shown are representative confocal images of actin-rich hair cell stereocilia (phalloidin, grey) at the cochlear base, mid, and apex of *Trim71^Δ/R595^* mice and control littermates. (J) Quantification of hair cells in (I) (n=3 animals per group, three independent experiments). Student *t* test was used to calculate the P values. \**P* < 0.05, \*\**P* < 0.01, \*\*\**P* < 0.001, and \*\*\*\**P* < 0.0001.

How far the differentiation of hair cells has advanced can be judged by the morphology of their actin-rich apical protrusions (termed stereocilia) and by the pattern of SGN innervation. Two distinct subtypes of SGNs innervate inner and outer hair cells (reviewed in ^32^). A single inner hair cell is innervated by multiple type I SGNs, whereas a single type II SGN, after a short turn towards the cochlear base, forms synapses with multiple outer hair cells. By E17.5/E18.5, inner and outer hair cells located at the base of the cochlea have formed stereocilia and outer hair cells are innervated by type II SGNs, a process that is guided by nearby Deiter’s cells and pillar cells^33^. Meanwhile, inner- and outer hair cells located at the apex lack stereocilia, and type II SGNs have yet to turn toward the base.

Our analysis of cochlear hair cell and SGNs morphology at stage E18.5 revealed that the innervation of outer hair cells at the base, mid, and apex was more advanced in *Trim7^R595H/Δ^* cochleae compared to control cochleae (Figure 4H). Furthermore, hair cells in *Trim7^R595H/Δ^* cochleae had a more mature stereocilia phenotype than hair cells in control cochleae (Figure 4I). Likely due to the shift in the timing of cell cycle exit and differentiation, we found that the hair cell density at the cochlear base and mid-turn was mildly reduced in E18.5 *Trim7^R595H/Δ^* mice compared to control littermates (Figure 4J). No changes in the timing of pro-sensory cell cycle exit and differentiation were observed in *Trim7^R595H/+^* mice (Figure S3).

### TGFβ-type signaling is upregulated in *Trim71* deficient cochlear progenitor cells

To investigate the molecular mechanisms leading to the premature differentiation of *Trim71*-deficient auditory pro-sensory cells, we administered dox at E8.5 and harvested cochlear epithelia from *Trim71^Δ/Δ^* mice and their control littermates at E13.5. We then isolated RNA for RT-qPCR and bulk RNA-sequencing (seq) experiments (Figure 5A). First, we validated conditional *Trim71* knockout using RT-qPCR. As expected, cochlear epithelia from E13.5 *Trim71^Δ/Δ^* mice lacked *Trim71* transcripts containing the critical exon4 but produced truncated, non-functional transcripts containing upstream exon2 and 3 (Figure 5B). Furthermore, consistent with early differentiation of hair cells, we found that the *Atoh1* transcript, which is expressed in nascent hair cells, was significantly upregulated in cochlear epithelia from E13.5 *Trim71^Δ/Δ^* mice compared to their control littermates (Figure 5C). Analysis of RNA-seq data identified 398 differentially expressed genes (DEG) (*p*-value <0.01) when comparing the transcriptome of *Trim71* deficient (*Trim71*^Δ*/*Δ^) cochlear epithelial cells to those of the controls (Figure 5D) (Table S1). Over three-quarters (340) of DEGs were upregulated *in Tim71* deficient cochlear epithelial cells, which aligns with TRIM71’s known inhibitory function in gene expression.

**Figure 5.**
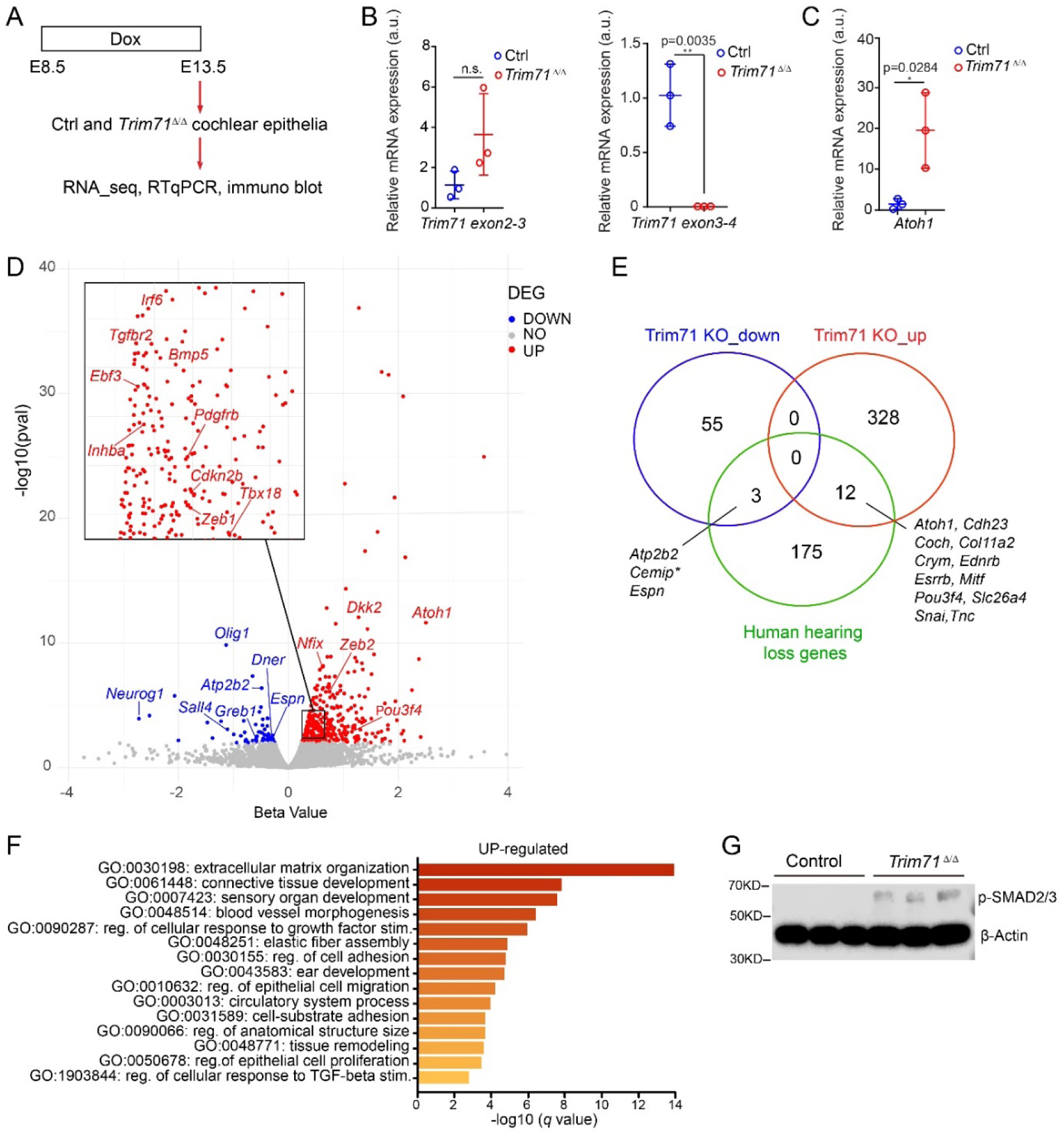
Loss of *Trim71* increases the expression of genes involved in sensory organ differentiation and extracellular matrix organization. (A) Experimental design: Cochlear epithelia were isolated from E13.5 *Trim71*^Δ*/*Δ^ embryos and *Trim71*^f/f^ (control) littermates following dox administration at E8.5. (B) RT-qPCR was used to validate the conditional ablation of *Trim71* (exon 4) and its associated transcripts in *Trim71*^Δ*/*Δ^ cochlear epithelia (n=3 animals per group, two independent experiments). (C) RT-qPCR reveals elevated *Atoh1* transcript levels in *Trim71*^Δ*/*Δ^ cochlear epithelia compared to control (n=3 animals per group, two independent experiments). (D) Volcano plot of RNA-seq data. Plotted is the beta-value (x-axis) versus −log10 q-value (y-axis). Transcripts that are significantly upregulated in *Trim71*^Δ*/*Δ^ cochlear epithelia are marked in red dots, and transcripts that are significantly downregulated are marked in blue dots. (E) Intersectional analysis to identify hearing loss genes that are altered by *Trim71* knockout. (F) Gene ontology enrichment analysis. Biological processes and pathways associated with genes that are upregulated in E13.5 *Trim71*^Δ*/*Δ^ cochlear epithelia are ranked by adjusted *P* value. (G) Immunoblots for p-SMAD2/3 and β-actin (loading control) using protein lysates of acutely isolated cochlear sensory epithelia from E13.5 *Trim71*^Δ*/*Δ^ embryos and control littermates (n=3, two independent experiments). Student *t* test was used to calculate the *P* values in (B) and (C). \**P* < 0.05, \*\**P* < 0.01, *P* >0.05 is deemed not significant (n.s).

We next investigated whether the expression of genes associated with hearing loss (HL) was affected (up or downregulated) by the loss of *Trim71*. We curated a list of 190 HL genes using the Walls WD, Azaiez H and Smith RJH Hereditary Hearing Loss Homepage (https://hereditaryhearingloss.org/) (Table S2), In total, we found that 15 HL genes were differentially expressed in Trim71 deficient cochlear epithelial cells compared to control, with three being downregulated and twelve being upregulated (Figure 5E).

To gain insights into the biological processes/ pathways that may be affected by the loss of *Trim71*, we conducted gene ontology enrichment analysis for up and downregulated genes ^34^. ‘Trans-synaptic signaling,’ ‘embryonic morphogenesis’, and ‘sensory perception of sound’ were among the biological processes that were significantly overrepresented in the list of downregulated genes (Table S3). Among the top 10 pathways/ biological processes that were significantly overrepresented in the list of upregulated genes were developmental processes such as ‘sensory organ’ and ‘ear development’ as well as processes associated with TGF-β-type signaling, including ‘extracellular matrix organization’, ‘connective tissue development’ and ‘regulation of cellular response to growth factor stimuli’ (Figure 5F) (Table S4) Notably, *Inhba,* which encodes for Activin A, a key ligand for Activin receptor signaling, and *Tgfbr2*, which encodes for TGF-β type II receptor (TGFBR2) were both upregulated. The activation of TGFβ type I and II receptors or Activin I and II receptors leads to the phosphorylation of effector proteins SMAD2 and SMAD3. After phosphorylation, these proteins translocate into the nucleus, where they bind to specific DNA binding elements and activate/repress transcription (reviewed in ^35^). To determine whether loss of *Trim71* leads to an over-activation of TGF-β type signaling, we analyzed phospho-SMAD2/3 levels in E13.5 cochlear epithelial protein lysates obtained from control and *Trim71^Δ/Δ^* embryos (n=3) (Figure 5G). We found that phospho-SMAD2/3 levels were higher in *Trim71* deficient cochlear epithelia than in control, suggesting that TRIM71 acts to repress premature activation of TGF-β-type signaling in cochlear epithelial cells.

### Disruption of Activin/TGF-β signaling delays cell cycle exit and differentiation of cochlear pro-sensory cells

We have recently shown that TGFβ2/TGFβ-RI signaling restricts cell cycle reentry and proliferation of early postnatal cochlear supporting cells in organoid culture ^36^. However, whether TGF-β signaling limits pro-sensory cell proliferation during cochlear development is currently unknown. To characterize the role of TGF-β signaling in the developing cochlea, we generated *Tgfbr1* conditional knockout (cKO) mice using *Pax2-Cre*, which is transiently expressed in otic progenitors ^37^. To determine the timing of terminal mitosis of pro-sensory cells, we administered a single pulse of EdU at E13.5 and analyzed EdU incorporation in hair cells and supporting cells (Deiter’s cells and pillar cells) at E18.5 in control *Tgfbr1* cKO mice. We found that loss of *Tgfbr1* mildly delayed pro-sensory cell cycle exit. We found that in *Tgfbr1* KO mice, the percentage of EdU-labeled hair cells (Figure S4A, B) and Deiter’s cells (supporting cell sub-type) (Figure S4A, D) in the cochlear apex was significantly higher than in control littermates. However, this mild delay in pro-sensory cell cycle exit did not increase the density of cochlear hair cells in the *Tgfbr1* deficient mice compared to control littermates (Figure S4C).

In previous research, we showed that the conditional knockout of *Inhba* using Pax2-Cre results in a mild delay in hair cell differentiation without affecting the timing of cell cycle withdrawal ^38^. To determine whether combined loss of *Tgfbr1* and *Inhba* exacerbates the defects in cell cycle withdrawal and differentiation observed in *Tgfbr1* and *Inhba* single knockout mice, we generated *Inhba* and *Tgfbr1* double knockout (DKO) mice using *Pax2-Cre*. EdU pulse-chase experiments, which involved a single EdU pulse at E13.5, revealed a significant increase in the number of EdU^+^ hair cells (Figure 6A, B) in the cochlear base and apex of E18.5 *Tgfbr1; Inhba* DKO mice compared to control. However, this mild delay in cell cycle exit did not significantly increase hair cell density in *Tgfbr1; Inhba* DKO mice compared to control littermates (Figure 6C). Furthermore, analysis of the outer hair cell innervation pattern (Figure 6D) and morphology of stereocilia (Figure 6E) revealed less mature phenotypes in *Tgfbr1; Inhba* DKO mice compared to control littermates. In sum, these findings suggest that *Trim71* maintains prosensory cells in a proliferative and undifferentiated state, at least in part by repressing TGF-β type signaling.

**Figure 6.**
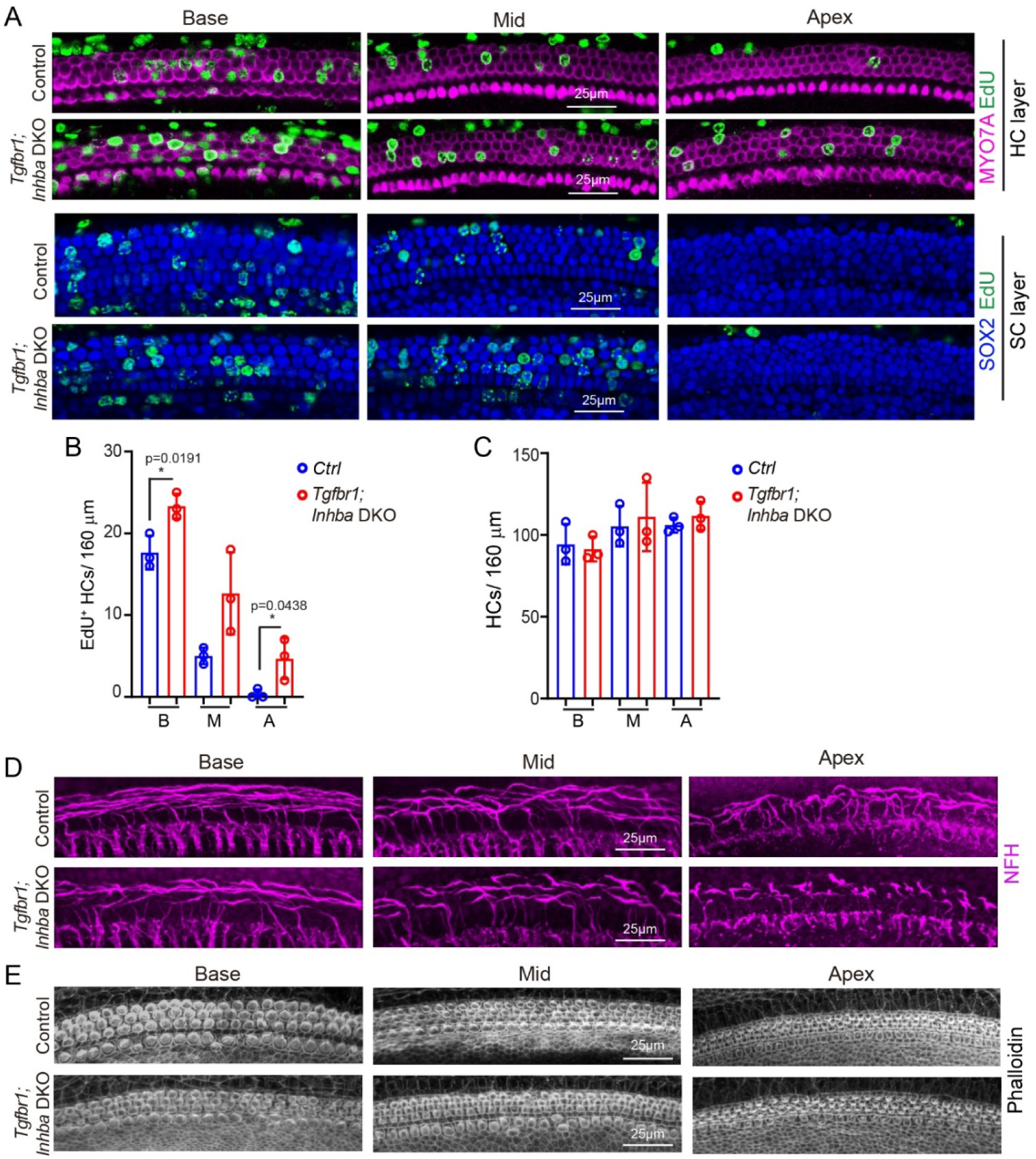
Loss of *Tgfbr1 and Inhba delays* cochlear pro-sensory cell *cycle exit* and differentiation. (A-E) Timed pregnant dams received EdU at E13.5, and cochlear tissue from *Tgfbr1-Inhba* double knockout mice (DKO) (*Pax2-Cre*; *Tgfbr1^f/f^*; *Inhba^f/f^*) and control littermates (*Tgfbr1^f/f^*; *Inhba^f/f^*) were collected and analyzed at E18.5. (A) Representative confocal images of EdU (green) incorporation in hair cells (MYO7A, magenta) and supporting cells (SOX2, blue) at cochlear apex, mid, and base in *Tgfbr1-Inhba* DKO mice and control littermates. (B) Cell cycle exit of hair cell progenitors is delayed in *Tgfbr1 Inhba* DKO mice. Quantification of EdU incorporation in hair cells in (A) (n=3 animals per group, three independent experiments). (C) Hair cell density is unchanged in *Tgfbr1 Inhba* DKO mice. Quantification of hair cell density in (A) (n=3 animals per group, three independent experiments). (D) Outer hair cell innervation is delayed in *Tgfbr1*; *Inhba* DKO mice. Confocal images of innervating neurons (NFH, magenta) at the cochlear base, mid, and apex of *Tgfbr1*; *Inhba* DKO mice and control littermates. (E) Hair cell stereocilia formation is delayed in *Tgfbr1*; *Inhba* DKO mice. Confocal images of hair cell stereocilia (phalloidin, grey) at the cochlear base, mid, and apex of *Tgfbr1*; *Inhba* DKO mice and control littermates. Student *t* test was used to calculate the *P* values in B and C.

### Loss of *Trim71* results in hair cell stereocilia and synaptic defects

HL-patients with mono-allelic missense mutations in *TRIM71* have, in addition to various middle and inner ear abnormalities, brain malformations (hydrocephaly). To determine whether brain malformations occur in our *Trim71* deficient mouse models, we imaged adult brains (P30) *Trim7^△/△^* and *Trim71^R595H/△^* mice and their control littermates using magnetic resonance imaging (MRI). Consistent with previous reports, we found that deletion of *Trim71* at E8.5 or thereafter does not cause hydrocephaly ^19^ (Figure S5A). To characterize inner ear/ middle ear morphology in adult (P30) *Trim7^△/△^* and *Trim71^R595H/△^* mice and their control littermates, we used Computed Tomography (CT) imaging. Our analysis revealed that *Trim7^△/△^* and *Trim71^R595H/△^* mice, like their control littermates (*Trim71*^+/+^ or *Trim71^R595H/+^*), had no obverse inner ear/middle ear malformations (Figure S5B). To further address the underlying mechanism of hearing loss, we analyzed whether hair cells degenerate in *Trim7^△/△^* and *Trim71^R595H/△^* mice or their stereocilia are lost or malformed. Our analysis revealed no reduction in the number of outer hair cells or inner hair cells in *Trim7^△/△^* and *Trim71^R595H/△^* mice compared to their control littermates (Figure S6A, C, D). However, closer examination of hair cell stereocilia revealed a modest but significant reduction in the length of inner hair cell stereocilia in *Trim7^△/△^* and *Trim71^R595H/△^* mice compared with *Trim71*^f/f^ (control) mice (Figure 7A, C). Further analysis of inner hair cell stereocilia length in *Trim7^△/△^* and control littermates revealed that the stereocilia length defect was limited to the apical and middle portion of the cochlea (Figure 7B, D).

**Figure 7.**
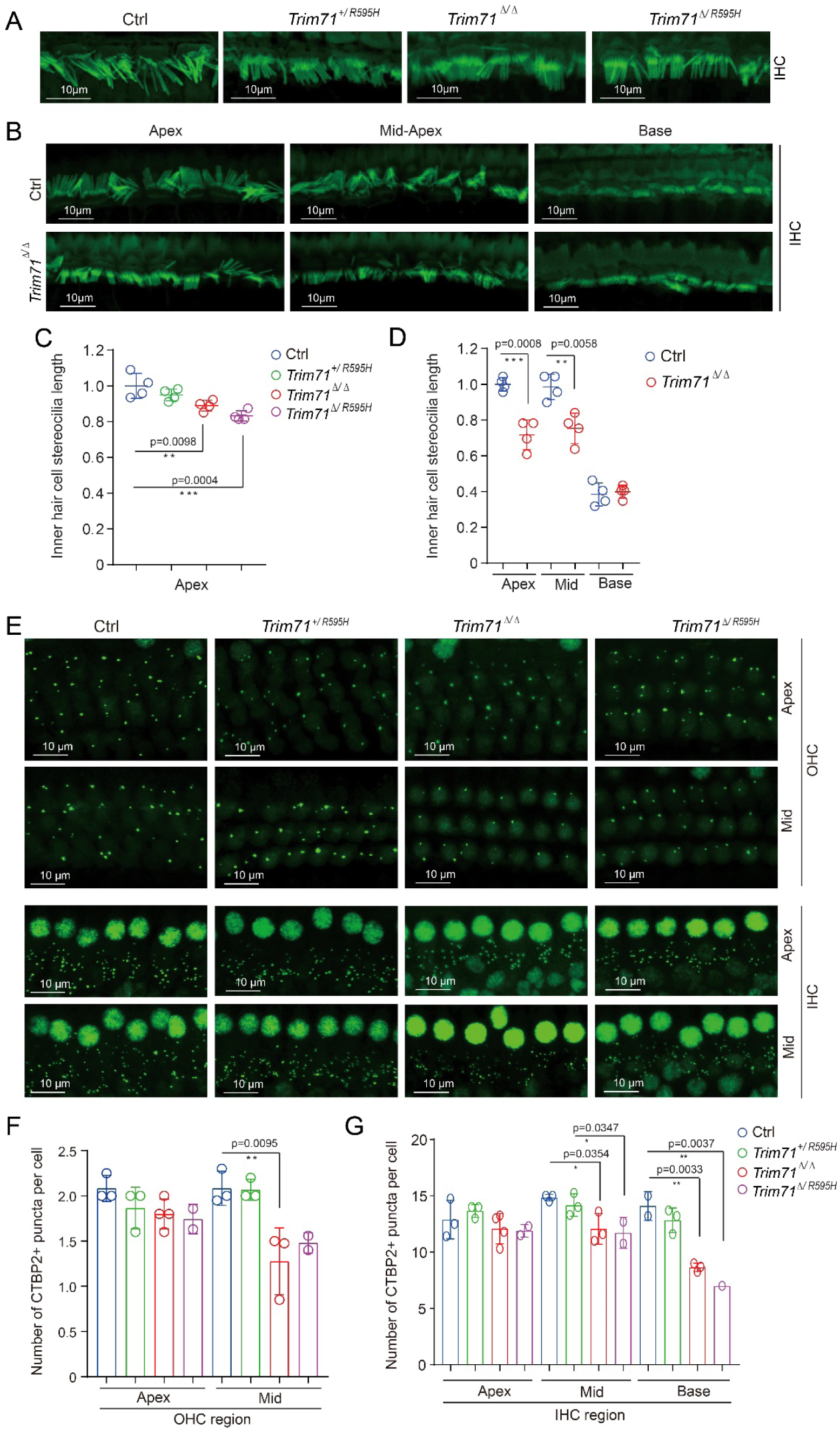
Loss of *Trim71* results in shorter inner hair cell stereocilia and fewer hair cell synapses. Dox was administered starting at E8.5, and *Trim71^Δ/Δ^* and *Trim71^R595H/Δ^* mice and their control littermates *Trim71^fl/fl^*(Ctrl), *Trim71^R595H/+^*were analyzed at P30. (A) Representative confocal images of inner hair cell stereocilia (phalloidin, green) in *Trim71^fl/fl^*(Ctrl), *Trim71^R595H/+^*, *Trim71^Δ/Δ,^* and *Trim71^R595H/Δ^* mice. Images were taken at the cochlear apex. (B) Representative confocal images of inner hair cell stereocilia (phalloidin, green) in *Trim71^fl/fl^*(Ctrl) and *Trim71^Δ/Δ^* mice. Images were taken at the cochlear apex, mid, and base. (C) Quantification of stereocilia length in A (n=4 animals per group). (D) Quantification of stereocilia length in B (n=4 animals per group). (E) Representative confocal images of pre-synaptic puncta (CTBP2, green) in outer hair cells (OHC) and inner hair cells (IHC). Images were taken at the cochlear apex, mid, and base. (F) Pre-synaptic puncta in outer hair cells are reduced in *Trim71^Δ/Δ^* mice. Quantification of CTBP2 puncta per outer hair cell (OHC) (n=3-4 for Ctrl, *Trim71^R595H/+^*, *Trim71^Δ/Δ^*, n=2 for *Trim71^R595H/Δ^*). (G) Pre-synaptic puncta in inner hair cells are reduced in *Trim71^Δ/Δ^* and *Trim71^R595H/Δ^* mice. Quantification of CTBP2 puncta per inner hair cell (n=3-4 for Ctrl, *Trim71^R595H/+^*, *Trim71^Δ/Δ^*, n=2 for *Trim71^R595H/Δ^*). One-way ANOVA was used to calculate the *P* values. *P* < 0.05, \*\**P* < 0.01 and \*\*\**P* < 0.001.

We next analyzed whether loss of *Trim71* alters the synaptic connections of hair cells with innervating type I and type II SGNs. To visualize hair cell-specific pre-synaptic complex, we used pre-synaptic marker protein CtBP2/RIBEYE ^39^. We found that the number of CtBP2 puncta per outer hair cells (Figure 7E, F) and inner hair cells (Figure 7E, G) was significantly reduced in the mid and basal portion of the cochlea but unchanged in cochlear apex in *Trim7^△/△^* and *Trim71^R595H/△^* mice compared to *Trim71*^f/f^ or *Trim71^R595H/+^* control littermates. Neurofilament H staining revealed a severe reduction in neuronal innervation of the outer hair cell region in *Trim7^△/△^* and *Trim71^R595H/△^* mice compared with *Trim71*^f/f^ or *Trim71^R595H/+^* control littermates (Figure S6A, B). Together, these data suggest that the loss of hearing observed in *Trim7^△/△^* and *Trim71^R595H/△^* mice may result from defects in synaptic transmission and or loss of neuronal modulation of outer hair cell function.

## DISCUSSION

Our study identifies *TRIM71* as a novel gene linked to hearing loss (HL). *TRIM71* missense mutations were first associated with a group of patients suffering from congenital hydrocephaly (CH) ^17^. Upon clinical reevaluation, it was determined that pathogenic variants in *TRIM71* are responsible for a syndromic phenotype that includes various features such as developmental delay, hearing loss, limb defects, craniofacial abnormalities, dysmorphism, and congenital cardiovascular defects with or without ventriculomegaly. Hereditary HL exhibits significant clinical and genetic heterogeneity (reviewed in ^40^). Hundreds of syndromic associations have been documented in which hearing loss coexists with abnormalities or malformations in other organs ^41^.

Most of the genes implicated in either isolated or syndromic HL code for proteins that regulate the development and or function of hair cells. In the case of syndromic HL, these proteins may also play roles in the development of semicircular canals, the middle ear, and the temporal bone (e.g. EYA1, SIX1, POU3F4, CHD7) ^42^. Some of these syndromes can present similar clinical features, and only genetic analysis can provide a definitive diagnosis. A clinical diagnosis of CHARGE syndrome was initially given to patient 1 due to the association of hearing loss with malformations of the external ear, middle ear, and lateral semicircular canals ^43^. Identifying a heterozygous pathogenic variant in *TRIM71* has allowed for the invalidation of this clinical diagnosis, leading to improved prognostic and genetic counseling for the patient.

Using *Trim71* knockout (*Trim71^Δ/Δ^*) and *Trim71* mutant mice that harbor the murine homolog of the human HL-associated missense mutation (*Trim71^R595H/Δ^*), we demonstrate a causative link between TRIM71 dysfunction and hearing loss. We find that loss of TRIM71 around∼ E9-E10, which correlates with the peak of murine *Trim71* expression in otic neural-sensory progenitors, impairs hearing, without causing structural inner ear or middle defects. *Trim71* homozygous mutant mice, which carry the R595H missense mutation and a conditional knockout allele, exhibited similar developmental defects as *Trim71* homozygous knockout mice. However, subtle differences in hearing sensitivity were observed. Specifically, *Trim71^R595H/Δ^* mice that received dox at E10.5 demonstrated high-frequency hearing loss, while the same timing of deletion in *Trim71^Δ/Δ^* mice did not affect hearing. The R595H miss-sense mutations in TRIM71 (R608H in humans) reside within TRIM71’s NHL domain, which previous studies have shown disrupts TRIM71’s RNA-binding ability without altering TRIM71’s ubiquitin ligase activity ^19^. Thus, the observed subtle differences in hearing sensitivity between *Trim71^R595H/Δ^* and *Trim71^Δ/Δ^* mice may be due to the preservation of TRIM71’s ubiquitin ligase activity in *Trim71^R595H/Δ^*mice. Only a small subset (<10%) of *Trim71* heterozygous mutant (*Trim71*^R595H/+^) mice manifest neuro-developmental defects ^19^. The only sporadic occurrence of neuro-developmental defects may explain why none of the *Trim71* heterozygous mutant (*Trim71*^R595H/+^, n=6) or heterozygous knockout mice (*Trim71*^Δ/+^, n=3) tested for hearing showed hearing deficits and suggests the existence of modifier gene(s).

The proper timing of auditory sensory development is disrupted by early otic (∼E9-E10) *Trim71* deletion in *Trim71^R595H/Δ^* and *Trim71^Δ/Δ^* mice. Cell cycle withdrawal and differentiation of auditory pro-sensory cells occur at precise times and follow opposing longitudinal gradients. Experiments conducted in murine and avian auditory organs have revealed that the RNA-binding protein LIN28B and members of *let-7* miRNA play a significant role in timing the exit from the cell cycle and differentiation of pro-sensory cells ^2,3^. However, the precise mechanism remained elusive. In this study, we demonstrate that TRIM71, a *let-7* target and member of the Lin28-let-7 pathway, is essential for preventing precocious pro-sensory cell cycle exit and differentiation. Our transcriptomic analysis revealed that cochlear progenitors in *Trim71^Δ/Δ^*mice expressed Activin A coding gene *Inhba* and TGFBRII coding gene *Tgfbr2* at significantly higher levels than its control counterpart. Our previous research has shown that Activin A acts as a pro-differentiation signal in the developing cochlea ^38^. However, while overexpression of Activin antagonist follistatin led to severe delay in pro-sensory cell cycle exit and hair cell differentiation, the differentiation defect in *Inhba* knockout mice was relatively mild. Also, we did not observe any defects in the timing of pro-sensory cell cycle exit in *Inhba* knockout mice, suggesting that other members of the TGFβ-family are compensating for the loss of *Inhba*. Indeed, our data from *Tgfbr1* single knockout and *Tgfbr1*-*Inhba* double knockout experiments indicate that TGF-β signaling cooperates with Activin signaling to drive pro-sensory cells out of the cell cycle and to promote their differentiation into hair cells and supporting cells. In addition to *Inhba* and *Tgfbr2*, several other known ‘pro-differentiation’ genes were upregulated in *Trim71* deficient cochlear progenitor cells. Notable among them are *Nfix* and *Ebf3*, which encode transcription factors that promote neurogenesis and gliogenesis ^44–46^. Future studies are warranted to address their function in cochlear sensory differentiation.

Besides the ear-related developmental defects, we found that *Trim71* knockout (*Trim71^Δ/Δ^*) mice and *Trim71* mutant (*Trim71^R595H/Δ^*) mice weighed less and had smaller bodies than *Trim71* heterozygous mutant/knockout or wild-type littermates. Most strikingly, both the *Trim71* knockout and *Trim71* mutant mice had unusually short tails. The small stature and the shortened tails resemble a phenotype reported for *let-7g* overexpressing mice, which lack caudal vertebrae due to a shortened developmental time window for axial elongation and somitogenesis ^47^.

How does early embryonic *Trim71* deficiency lead to hearing loss in mice? In humans, *Trim71* missense mutations lead to both sensorineural and conductive hearing loss, which is associated with malformations of the middle ear and temporal bone. However, MRI and CT imaging of the inner ear temporal bone and middle ear bones of adult *Trim71^R595H/Δ^* or *Trim71^Δ/Δ^* mice (dox E8.5) showed no defects. This suggests that the observed hearing loss in *Trim71* homozygous mutant/knockout mice is due to the dysfunction of hair cells and or neurons. While the precise mechanism remains unresolved, our transcriptomic and morphological analysis have identified several candidates. We found that in *Trim71* deficient cochlear epithelial progenitors, the expression of mesenchymal transcription factors (e.g., *Pou3f4, Snai2, and Zeb2*) is upregulated. The vital role of otic mesenchyme for cochlear development and hearing is best exemplified by the mesenchymal-specific transcription factor POU3F4. POU3F4 regulates temporal bone development ^48^, but also spiral ganglion fasciculation ^49^ and mutations in *POU3F4* (DFNX2) are the leading cause of non-syndromic X-linked hearing loss in humans ^50,51^. Future studies will investigate whether and to what abnormal expression of mesenchymal transcription factors in cochlear epithelial progenitors leads to hearing loss.

Furthermore, we found that two well-known HL genes, *Espn* (DFNB36) ^52^ and *Atp2b2* (DFNA82) ^53^, were downregulated in *Trim71* deficient cochlear progenitor cells. Both ATP2B2 (also referred to as PMCA2) and ESPN proteins localize to hair cell stereocilia, where they regulate Ca2+ homeostasis ^54^ and stereocilia morphology ^55^ respectively. Our examination of stereocilia morphology in adult *Trim71* homozygous mutant/ knockout mice revealed a modest but significant reduction in the length of inner hair cell stereocilia in the apex and mid portion of the cochlea compared to control littermates. Stereocilia shortening is also observed when mechano-transduction is disrupted ^56^, and future studies will address whether or not mechano-transduction is altered *Trim71* homozygous mutant/ knockout mice. Synaptic transmission and signal amplification defects are other possible causes for the loss of hearing observed in *Trim71* homozygous mutant/ knockout mice. Specifically, we found that inner and outer hair cells in *Trim71* homozygous mutant/ knockout mice had fewer pre-synaptic puncta than control littermates. Additionally, in a subset of *Trim71* homozygous mutant/ knockout mice, neuronal innervation of the outer hair cell region-which is essential for signal amplification and modulation-was nearly absent. It is intriguing to consider that the changes in the timing of terminal mitosis and the differentiation of hair cell and supporting cell progenitors may have led to the structural changes observed in *Trim71* homozygous mutant/ knockout mice. Alternatively, TRIM711 might serve additional functions during early otic development that are not related to its influence on developmental timing. Future studies are needed to investigate TRIM71’s role in the early stages of otic development.

## MATERIALS AND METHODS

### Patient data

Whole-exome sequencing was performed on three members of patient 1’ family, using the method previously described ^57,58^. Patient 2’s genetic testing has been recently described^18^. Patient 1 and patient 2 were clinically examined by an Ear Nose and Throat physician and a clinical geneticist.

They had air and bone conduction pure tone audiograms, and hearing impairment was defined according to the recommendations of the International Bureau for Audiophonology (BIAP) https://www.biap.org/en/recommandations/recommendations/tc-02-classification.

MRI and CT scans of the internal auditory canal, labyrinth, and middle ear were performed and analyzed using methods previously described ^59,60^.

This study was approved by the Institutional Review Board at Massachusetts General Hospital and Yale University. All genetic testing was obtained with written informed consent prior to collection from patients.

### Mouse breeding and genotyping

All experiments and procedures were approved by the Johns Hopkins University Institutional Animal Care and Use Committees protocol, and all experiments and procedures adhered to National Institutes of Health-approved standards. *Trim71^R595H^* mutant mice (RRID: MGI:7661051) were provided by Kris Kahle, Harvard Medical School. *Trim71* floxed mice (RRID: MGI:7661049) were provided by Waldemar Kolanus, Universität Bonn. *Pax2-Cre* transgenic mice (RRID: MGI:3046196) were obtained from Andrew Groves, Baylor College. *Inhba* floxed mice (RRID: MGI:3758877) were obtained from Martin Matzuk, Baylor College. *Tgfbr1* floxed (RRID: IMSR_JAX:028701), *TetO-cre* (RRID: IMSR_JAX:006234) and *R26^rtTA*M2^* (RRID: IMSR_JAX:006965) mice were purchased from Jackson Laboratories (Bar Harbor, ME). To induce Cre expression, doxycycline (dox) was delivered to time-mated females via ad libitum access to feed containing 2g/kg dox. Mice were genotyped by PCR as previously published. Genotyping primers are listed in Table S5. Mice of both sexes were used in this study. Embryonic development was considered as E0.5 on the day a mating plug was observed. All animal work was performed in accordance with approved animal protocols from the Institutional Animal Care and Use Committees at Johns Hopkins University School of Medicine.

### RNA extraction and RT-qPCR

Organoids were harvested using Cell Recovery Solution (Corning, no.354253). Total RNA from organoids/tissue was extracted using the miRNeasy Micro Kit (QIAGEN, no. 217084). MRNA was reverse transcribed into cDNA using the iScript cDNA synthesis kit (Bio-Rad, no. 1708890). Q-PCR was performed on a CFX-Connect Real Time PCR Detection System using SYBR Green Master Mix reagent (Thermo Fisher Scientific, no. 4385612). Gene-specific primers used are listed in Table S6. *Rpl19* was used as the endogenous reference transcript. Relative gene expression was calculated using ^ΔΔ^CT method ^61^.

### RNA sequencing and data analysis

To induce *Trim71* deletion, a timed pregnant female received dox-containing feed starting at E8.5. Offspring were collected at E13.5, and cochlear epithelial ducts from individual *Trim71* KO (*Trim71* ^Δ/Δ^) and control (*Trim71*^f/f^) animals were isolated. After RNA extraction, samples were processed using Illumina’s TruSeq stranded Total RNA kit, per the manufacturer’s recommendations. The samples were sequenced on the NovaSeq 6000, paired-end, 2x50 base pair reads. Kallisto (v0.46.1) ^62^ was used to pseudo-align reads to the reference mouse transcriptome and to quantify transcript abundance. The transcriptome index was built using the Ensembl Mus Musculus v96 transcriptome. The companion analysis tool Sleuth was used to identify differentially expressed genes (DEGs)^63^. These lists were then represented graphically using sleuth, pheatmap and ggplot2 packages in R v1.3.1093. Gene identifier conversion, gene annotation, and enrichment analysis were conducted using Metascape ^34^.

### Immunohistochemistry

Cochleae were fixed with 4% (vol/vol) paraformaldehyde in PBS (Electron Microscopy Sciences, no. 15713) overnight at 4 degrees. The P30 cochlea tissue was decalcified in 5% EDTA for 2 days following fixation. To permeabilize cells and block unspecific antibody binding, organoids/explants were incubated with 0.5% (vol/vol) TritonX-100/10% (vol/vol) FBS in 1× PBS for 30 min. Antibody labeling was performed according to manufacturer’s recommendations. Antibody information is listed in Table S7.

### Cell Proliferation

EdU (25mg/kg, E10187, Thermo Fisher Scientific) was intraperitoneal injected on embryonic day E13.5 and cochleae were harvested at E18.5. EdU incorporation was detected using Click-iT Edu Alexa Fluor 555 imaging Kit (Thermo Fisher Scientific, no. C10338) following the manufacturer’s specifications.

### Immunoblotting

Cochlear epithelia cells from E13.5 were lysed with RIPA buffer (Sigma-Aldrich, no. R0278) supplemented with protease inhibitor (Sigma-Aldrich, no.11697498001), phosphatase Inhibitor mixture 2 (Sigma-Aldrich, no. P5726) and phosphatase inhibitor mixture 3 (Sigma-Aldrich, no. P0044). Western blots were generated as described previously ^64^. Antibody information is listed in Table S7.

### Auditory brainstem response (ABR) measurements

ABR measurements were performed following published procedures ^65^. Briefly, 4–5 week– old mice were anesthetized with Ketamine (100 mg/kg) and Xylazine (20 mg/kg) and placed on a heating pad inside a sound-attenuating chamber. Subdermal platinum needle electrodes (E2, GrassTechnologies, West Warwick, RI) were placed on the left pinna (inverting), vertex (non-inverting), and the leg muscle (ground). ABR stimuli were delivered through a free-field speaker (FD28D, Fostex, Tokyo, Japan) placed 10 cm away from the animal’s head. ABR stimuli generation and signal acquisition were controlled by a BiosigRz software interfacing TDT WS4 high-performance computer workstation. ABRs, pure tone bursts at 4, 8, 16, 24, and 32 kHz were presented at a rate of 21/s, with a 5 ms duration. Stimuli were presented in descending sound levels from the maximum speaker output level in 10dB increments. Responses were collected using a low impedance head stage (RA4L1; TDT), pre-amplified and digitized (RA4PA preamp; TDT), and sent to an RZ6 processing module. The signal was filtered (300-3000 Hz) and averaged over 512 presentations. The sound threshold was defined as the sound level at which the peak-to-peak ABR signal magnitude in wave I was two standard deviations above the average background noise level.

### Magnetic resonance imaging (MRI) and computer tomography (CT)

Mouse heads were fixed with 4% (vol/vol) paraformaldehyde in PBS overnight and then mouse structural CT imaging data were collected with TriFoil Small Animal Tomography Systems in the core facility of Johns Hopkins University. Mouse structural MR imaging data were collected with Bruker 11.7T in the core facility of Johns Hopkins University.

### Statistical Analysis

All results were confirmed by at least two independent experiments. The sample size (n) represents the number of animals analyzed per group. Animals (biological replicates) were allocated into control or experimental groups based on genotype and/or type of treatment. To avoid bias, masking was used during data analysis. Data was analyzed using GraphPad Prism 8.0. Relevant information for each experiment, including sample size, statistical tests, and reported p-values, are found in the legend corresponding to each figure. In all cases, p-values ≤0.05 were considered significant and error bars represent standard deviation (SD).

### Data and materials availability

RNA sequencing data have been deposited in the Gene Expression Omnibus data repository under accession GSE281437

## ACKNOWLEDGMENTS

We thank the members of the Doetzlhofer Laboratory for the help and advice provided throughout the course of this study. The authors are grateful to patients, Fondation pour l’audition, the Association « S’entendre », and the Centre de Ressources Biologiques of Imagine Institute.

## Author Contributions

Conceptualization: A.D., X.-J.L, K.K., S.M.

Methodology: A.D., X.-J.L, L.E., PT.NP, C.M., W.K., P.D, S.M.

Investigation: X.-J.L., L.E., PT.NP., C.M, S.M., R.M., K.B., S.E., F.D., C.M., F.B.

Supervision: A.D., K.K., S.M.

Writing—original draft: A.D., X.-J.L, S.M.

Writing—review& editing: A.D., X.-J.L., C.M, W.K., P.D., K.K, S.M.

## Supplemental information (SI)

**SI Appendix.** Figure S1-Figure S6, Table S5-S7

**Tables S1-S4** are provided as separate files in spreadsheet format (excel file)

**Table S1** Wald test Ctrl vs. *Trim71-KO*

**Table S2** Hearing loss gene list

**Table S3** GO analysis of downregulated genes Ctrl vs. *Trim71*-KO

**Table S4** GO analysis of upregulated genes Ctrl vs. *Trim71-KO*

