## Supplementary information for "The RNA-binding protein TRIM71 is essential for hearing in humans and mice and regulates the timing of auditory sensory organ development"

**Supplemental information**

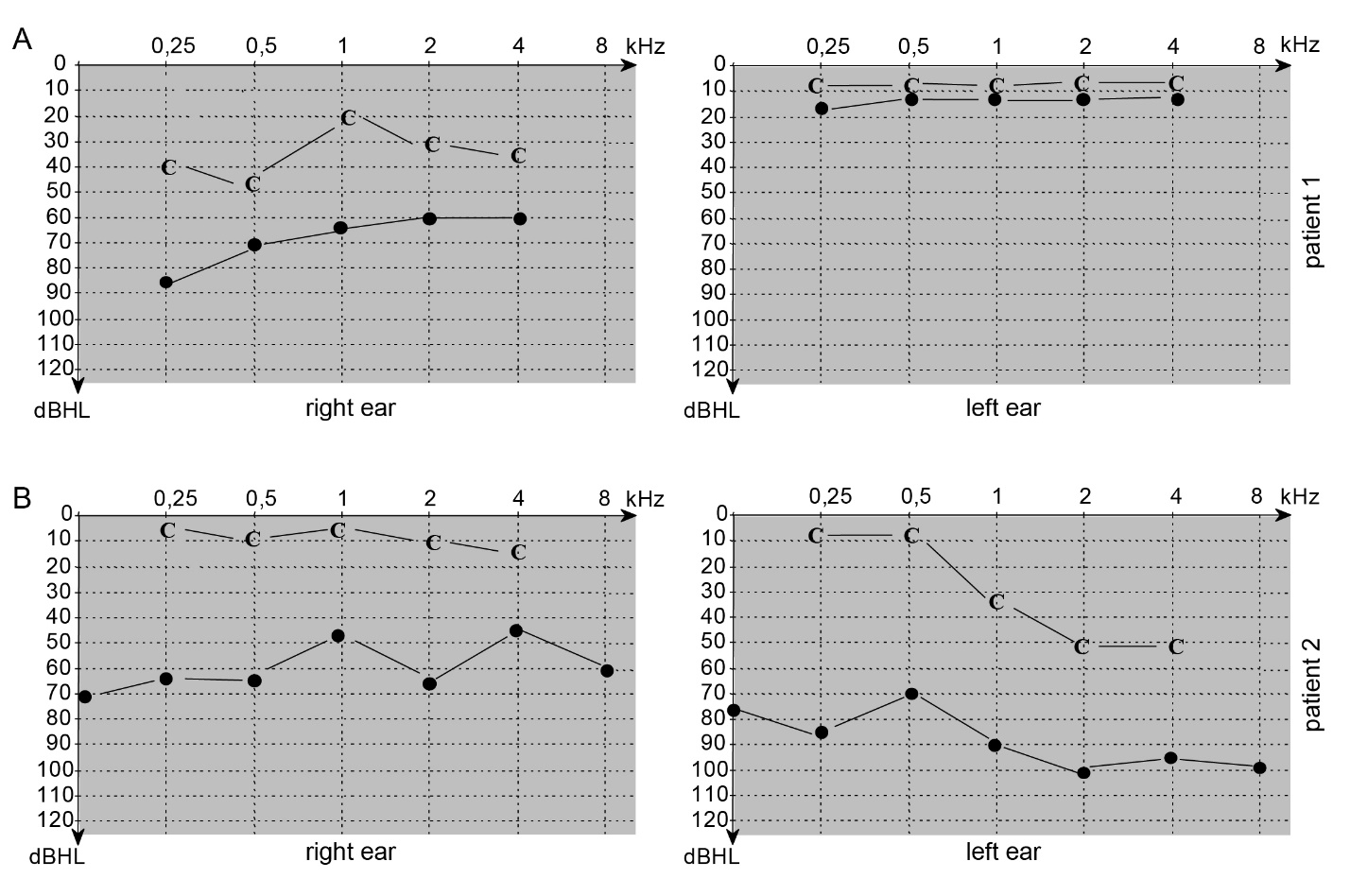

**Figure S1 CH patients with a missense mutation in *TRIM71* exhibit mixed hearing impairment**. Patient 1 has a missense mutation of W334R, and patient 2 has a missense mutation of R608H. (A-B) Audiograms. Plotted are on the x-axis the sound frequency in kHz and on the y-axis sound intensity in decibels hearing level (dBHL). Points = air conduction thresholds; C = bone conduction thresholds. (A) Patient 1’s audiogram at 5 years old identified a mixed (conductive and sensorineural) severe right hearing impairment. (B) Patient 2’s audiogram at 6 years old identified a mixed (conductive and sensorineural) severe left and a moderate conductive right hearing impairment.

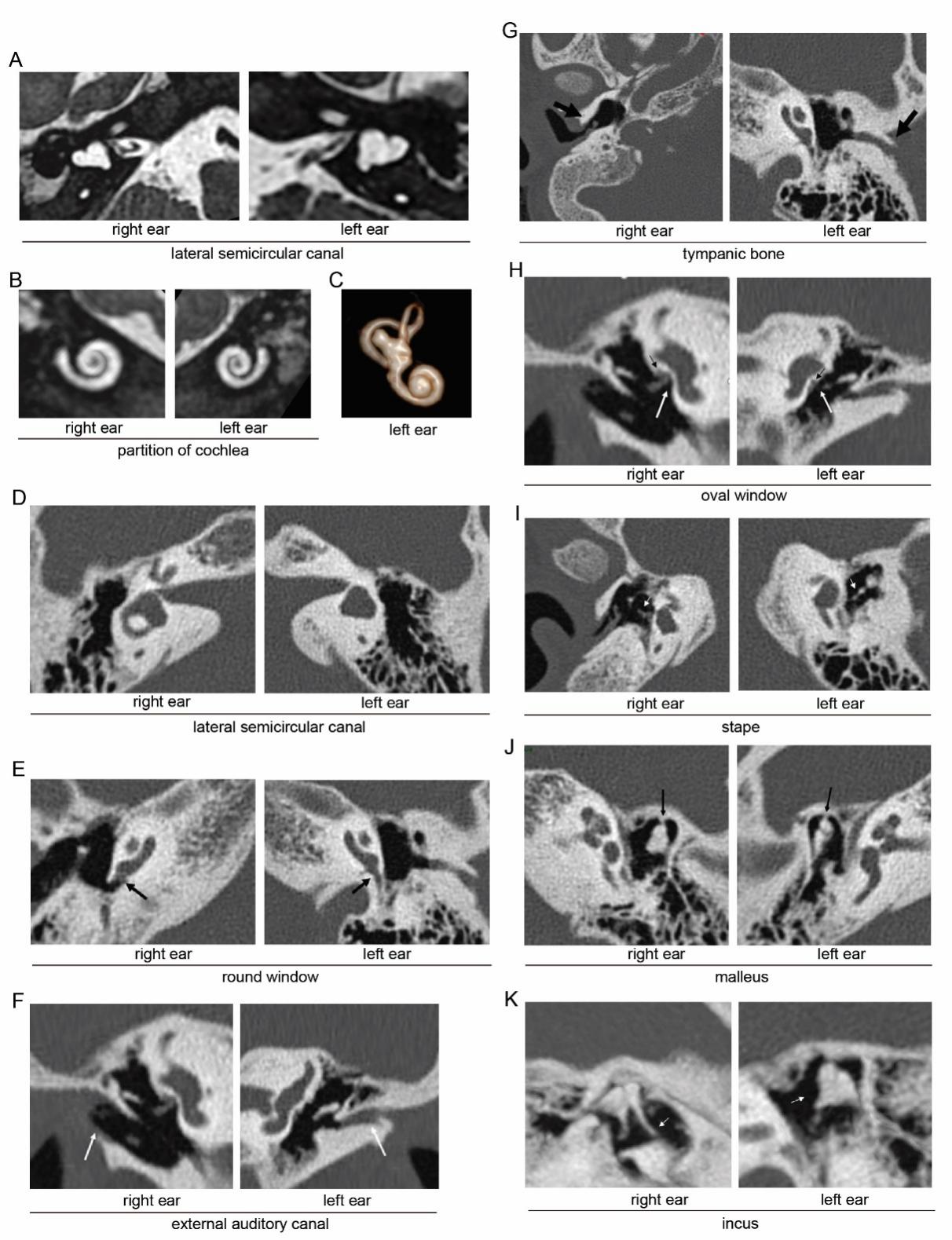

**Figure S2 CH patients with a missense mutation in *TRIM71* exhibit inner ear and middle ear malformations.** (A-C) Patient’s 1 inner ear MRI. (A) MRI 3DT2HR sequence axial reconstruction in the plane of the lateral semicircular canal of the right and left labyrinth: vesicular and short bilateral appearance of the lateral semicircular canal. (B) MRI 3DT2HR sequence coronal MIP (Maximum Intensity Projection) reconstruction in the plane of the basal cochlea of the right and left labyrinth: normal appearance of the partition of the right and left cochlea. (C) MRI Volume rendering 3DT2 axial reconstruction in the plane of the left osseous labyrinth vesicular and short appearance of the lateral semicircular canal. (D-K) Patient 2’s temporal bone CT scan. (D) Axial CT reconstruction of the left and right temporal bone on the plane of the lateral semicircular canal. Normal appearance of the right lateral semicircular canal and short appearance of the left lateral semicircular canal with an absence of central bone island.

(E) Normal appearance of the partition of the right and left cochlea; note the narrowness of the left round window and the normal right window (black arrow). (F-G): Coronal (F) and axial (G) CT reconstruction of the right and left temporal bone: stenotic left external ear canal and small but present right external auditory canal (F, white arrow). Note the thick and dysplastic appearance of the right tympanic bone and the hypoplastic left tympanic bone (G, black arrow). (H-I) Coronal (H) and axial (I) CT reconstruction of the right and left temporal bone in the plane of the stapes: narrowness of the right oval window and atretic left oval window (H, white arrow), bilateral aberrant course of the facial nerve's tympanic segment next to the oval window (H, black arrows). Note the monopod left stapes and the dysplastic stocky right stapes (I, white arrow). (J-K): Axial oblique CT reconstruction in the plane of the malleus and incus. J: bilateral fixation of the head of the malleus (black arrows). K: normal appearance of the malleus and long process of the incus in the right ear and fusion of the head of the malleus with the body of the incus and aplasia of the handle of the malleus and the long process of the incus in the left ear (white arrow).

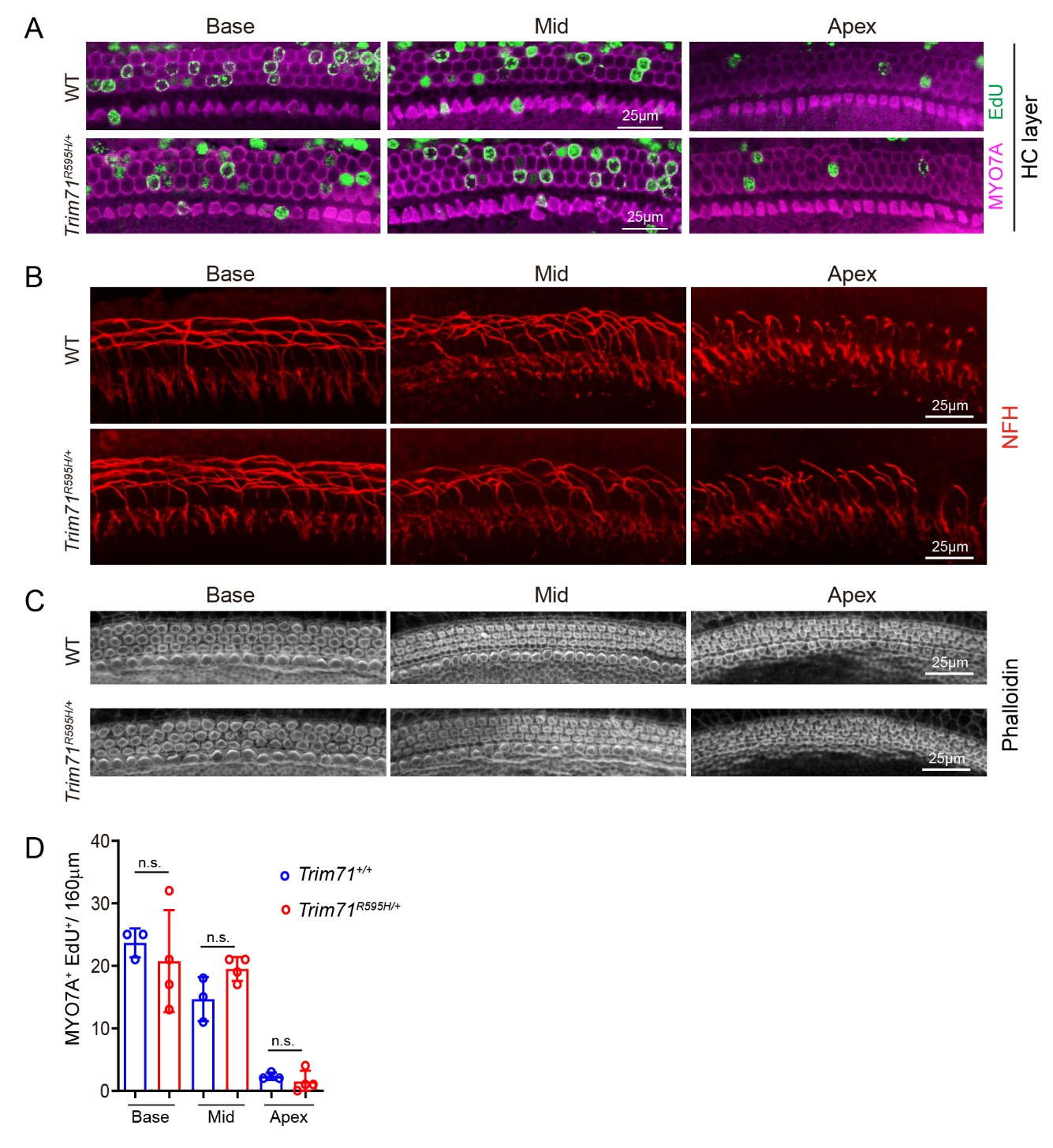

**Figure S3. Cochlear pro-sensory cell cycle withdrawal and differentiation are normal in *Trim71^R595H/+^* mice.** (A) Representative confocal images of hair cell (HC) layer of cochlear sensory epithelia of E18.5 *Trim71^+/+^* (wild type, WT) and *Trim71^R595H/+^* mice. EdU incorporation (green) in hair cells (MYO7a, magenta) was analyzed following a single EdU pulse at E13.5. (B) Neurofilament-heavy chain staining (NFH, red) was used to analyze outer hair cell innervation patterns in stage E18.5 *Trim71^R595H/+^* mice and *Trim71^/+^* (control) littermates. (C) Phalloidin staining was used to visualize hair cells' stereocilia morphology at the cochlear base, mid, and apex from E18.5 *Trim7^+/+^* (control) and *Trim71^R595H/+^* embryos. (D) Quantification of EdU-positive MYO7A^+^ hair cells in (A) (n=3 in WT mice and n=4 in *Trim71^R595H/+^*, two independent experiments).

A two-tailed, unpaired Student’s t-test was used to calculate *P* values. *P* >0.05 was deemed not significant (n.s.).

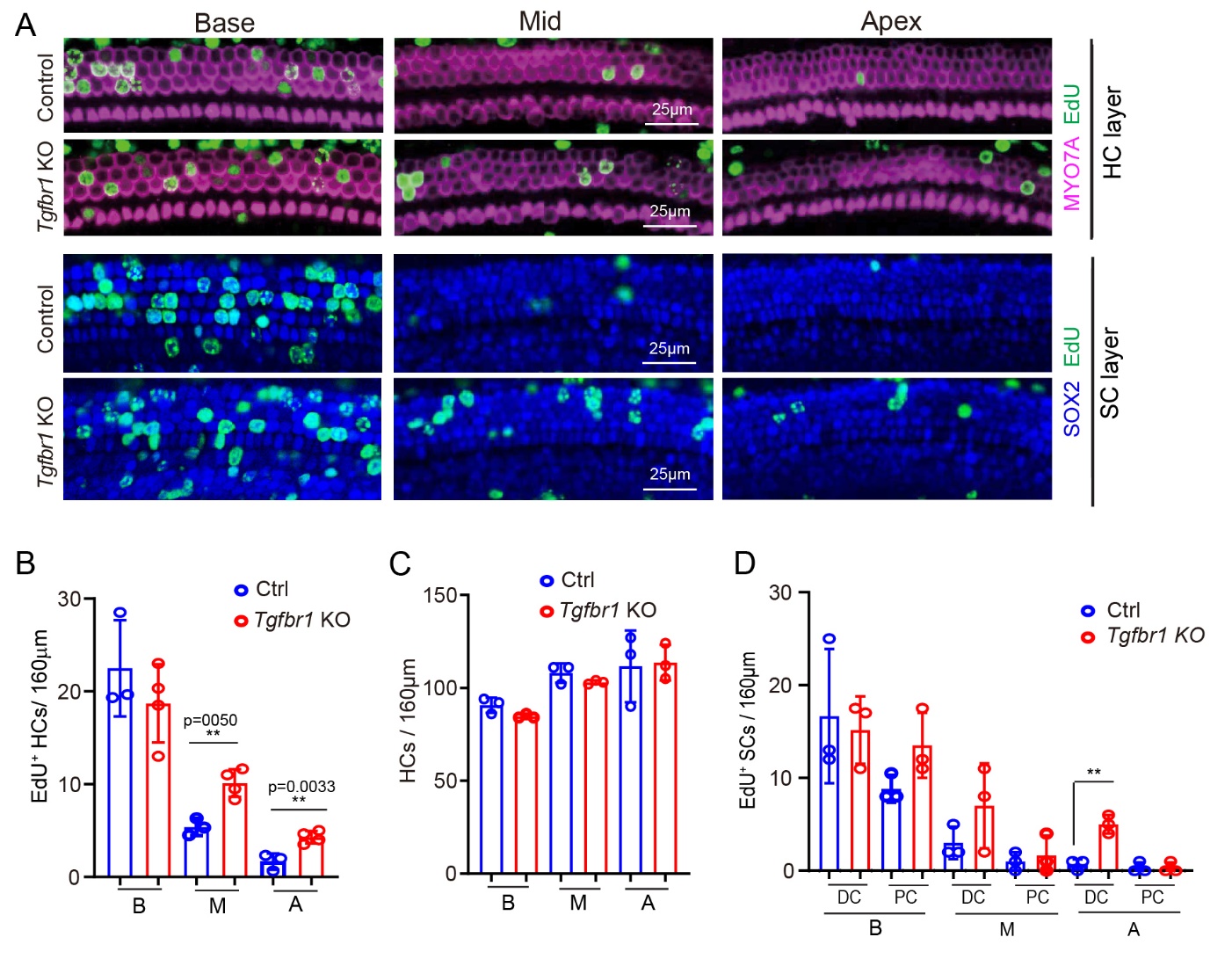

**Figure S4. Deletion of *Tgfbr1* delays cell cycle exit of auditory pro-sensory cells.** (A-D) Timed pregnant dams received EdU at E13.5, and EdU incorporation in hair cells (HCs) and supporting cells (SCs) was analyzed at E18.5 in *Tgfbr1* KO (*Pax2-Cre; Tgfbr1^f/f^)* mice and control (*Tgfbr1 ^f/f^*) littermates. (A) Representative confocal images of apical, mid, and basal segments of cochlear sensory epithelia of *Tgfbr1* KO mice *and* control littermates. Images show EdU (green) incorporation in hair cells (MYO7A, magenta) and supporting cells (SOX2, blue). (B) Cell cycle exit of hair cell progenitors is delayed in *Tgfbr1* KO mice. Quantification of EdU-positive hair cells (HCs) in (A) (n=3, three independent experiments). (C) Hair cell numbers are unchanged in *Tgfbr1* KO mice. Quantification of hair cell density in base, mid, and apex (HCs) in (A) (n=3, three independent experiments). (D) Cell cycle exit of supporting cell progenitors is delayed in *Tgfbr1* KO mice EdU incorporation in supporting cells (SCs) in (A) (n=3, three independent experiments). Two-tailed, unpaired Student’s *t* test was used to calculate *P* values. **P* ≤ 0.05, ***P* <0.01.

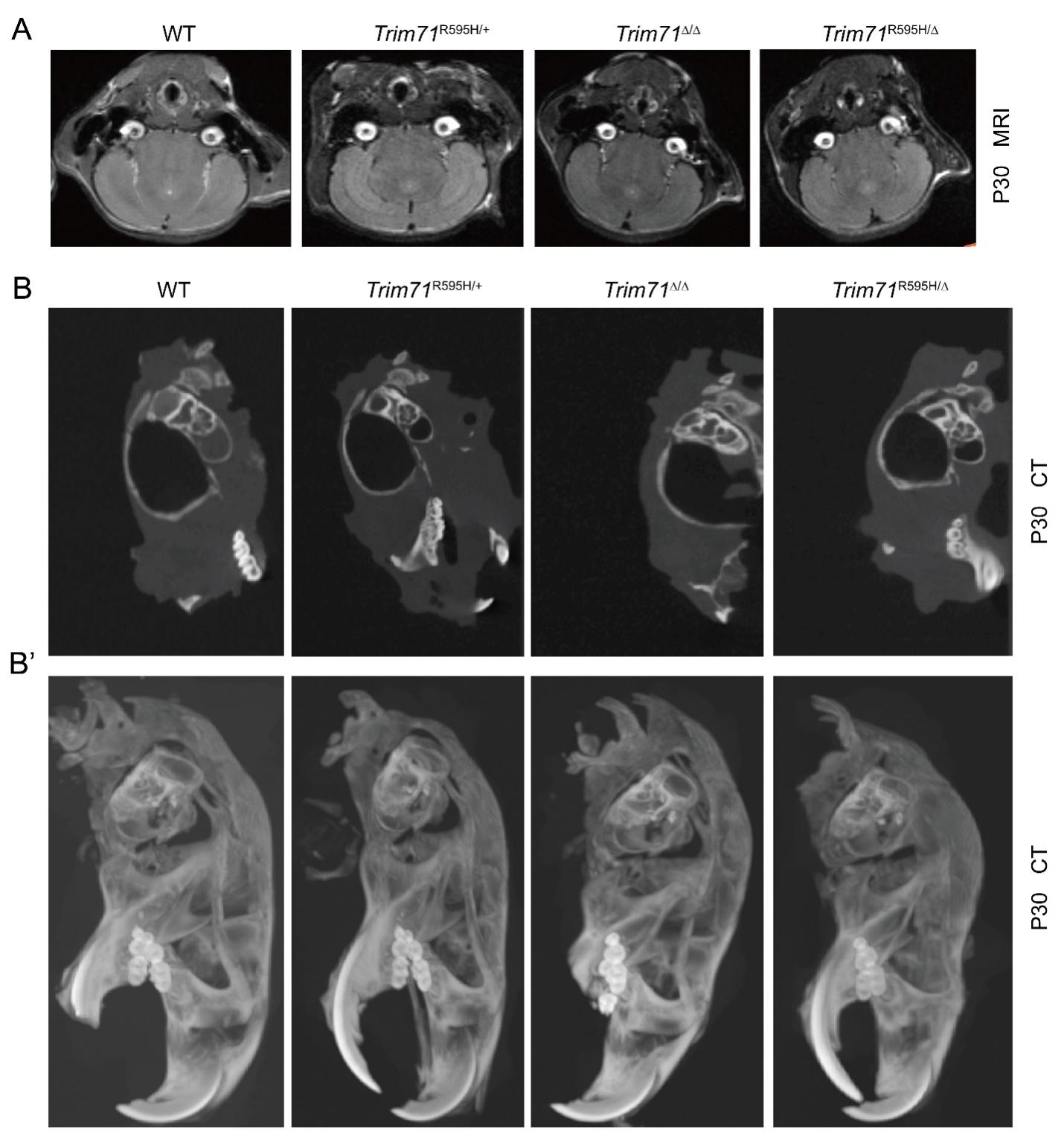

**Figure S5.** Head CT and MRI scans of stage P30 *Trim71^Δ/Δ^* and *Trim71^R595H/Δ^* mice and their control littermates (*Trim71^fl/fl^*, *Trim71^R595H/+^*). Dox was administered at E8.5. (A) MRIs show that *Trim71^Δ/Δ^* and *Trim71^R595H/Δ^* mice have no brain abnormalities. (B) CT scans reveal that *Trim71^Δ/Δ^* and *Trim71^R595H/Δ^* mice have normal inner ear and middle ear anatomy. (B’) Whole volume CT reconstruction.

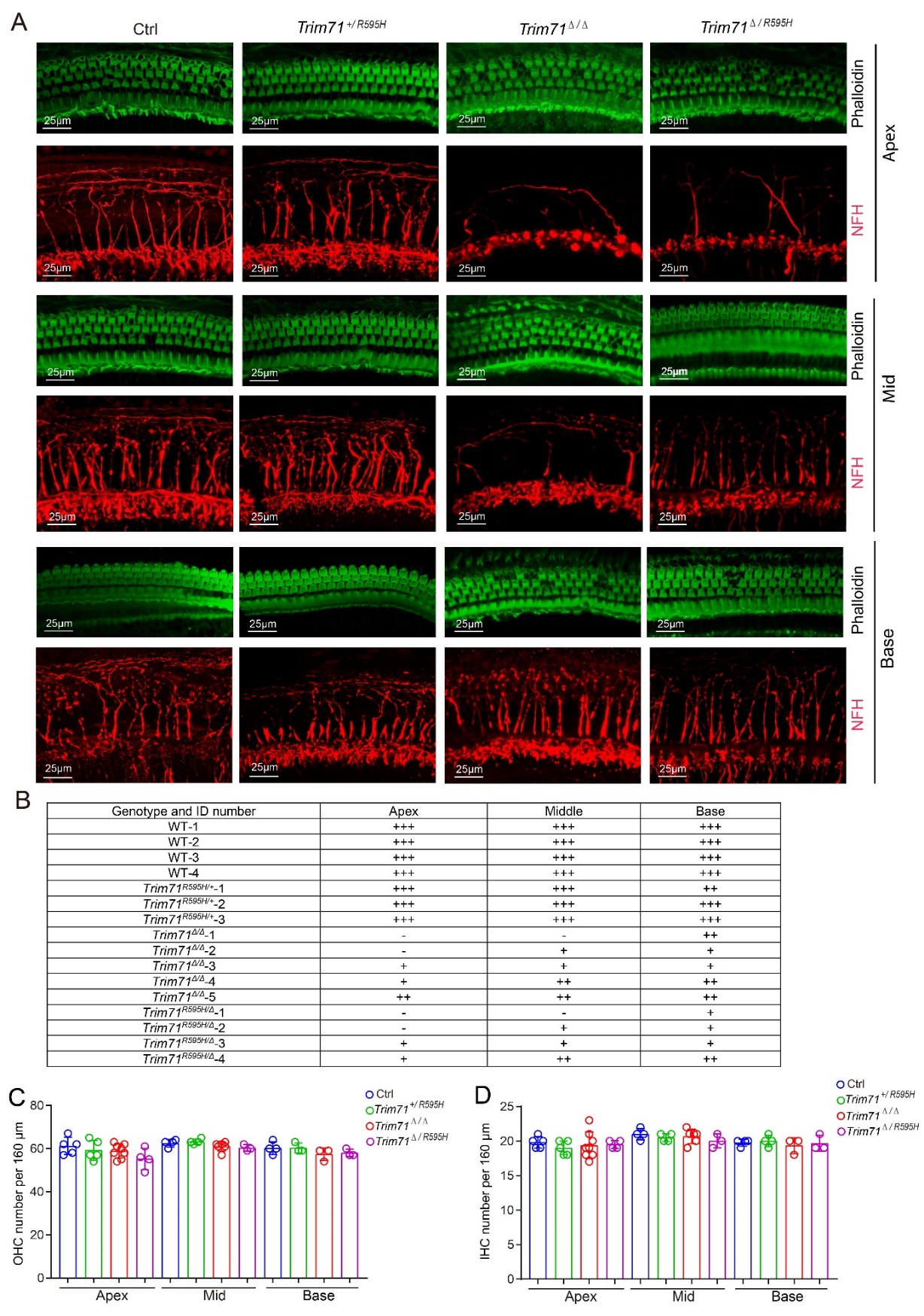

**Figure S6. Early otic deletion of Trim71 causes neuronal degeneration.** To induce conditional deletion of Trim71 floxed allele(s), timed-mated pregnant dames received doxycycline-containing feed starting at E8.5, and offspring were analyzed at P30. (A) Low power confocal images of auditory sensory epithelium in 1-month-old *Trim71^fl/fl^* (WT), *Trim71^R595H/+^*, *Trim71^Δ/Δ^* and *Trim71^R595H/Δ^* mice. Phalloidin staining (green) labels inner and outer hair cell stereocilia, and neurofilament H (NFH, red) immuno-staining labels innervating neuronal fibers. Shown are representative images of the cochlear apex, mid, and base. (B) Table summarizing neuronal innervation of outer hair cell region in 1-month-old *Trim71^fl/fl^* (WT), *Trim71^R595H/+^*, *Trim71^Δ/Δ^*, and *Trim71^R595H/Δ^* mice. The following scores indicate: (+++) no innervation defect, (++) mild defect, (+) severe defect, (-) innervation completely missing. (C-D) Outer hair cell (OHC) density (C) and inner hair cell (IHC) density (D) in the cochlear apex, mid and base of control (*Trim71^fl/fl^*), *Trim71^R595H/+^*, *Trim71^Δ/Δ^*, and *Trim71^R595H/Δ^* mice. Graphed are individual data points representing average values per animal (n = 3-8 per group).

**Table S5 Genotyping primers.**

| **Mouse line** | **Genotyping primers** | **Product size** |
| --- | --- | --- |
| *R26 ^rtTA*M2^* | MTR: GCG AAG AGT TTG TCC TCA ACC  F: AAA GTC GCT CTG AGT TGT TAT  WTR: GGA GCG GGA GAA ATG GAT ATG | WT=650bp  MT=340bp |
| *Trim71 floxed* | Trim71-F-WT: GAA AGG AGG CTA GCC AAA GG  Trim71-R-KO: ATG CTG TAC GGT AGG AGT CTT CC | FL=350bp  WT=250bp |
| *Trim71^R595H/+^* | F: GAC GGA AAC CTG TTT GGT GC  R: GTC GGC CAC TAT GAT CCT GC | WT=420bp  Mutant=229 and 192bp |
| *TetO-Cre and Pax2-Cre* | F: GCC TGC ATT ACC GGT CGA TGC AAC GA  R: GTG GCA GAT GGC GCG GCA ACA CCA TT | TG:700bp |
| *Inhba floxed* | Inhba-fx-1: AAG AGA GAA TGG TGT ACC TTC ATT  Inhba-fx-2: TAT AAC CTG GGT AAG TGG GT  Inhba-fx-3: AGA CGT GCT ACT TCC ATT TG | FL=400bp  WT=280bp |
| *Tgfbr1 floxed* | F: CCT GCA GTA AAC TTG GAA TAA GAA G  R: GAC CAT CAG CTG TCA GTA CCC | WT=219bp  Mutant=320 |

**Table S6 QPCR primers.**

| **Gene** | **Forward Primer** | **Reveres Primer** |
| --- | --- | --- |
| *Atoh1* | ATG CAC GGG CTG AAC CA | TCG TTG TTG AAG GAC GGG ATA |
| *Hmga2* | CAG AAG AAA GCA GAG ACC ATT GG | TTG TTG TGG CCA TTT CCT AGG T |
| *Trim71(Exon3/4)* | ATC GGG AGT GTG AGC TGT TG | GGC GTG AAC ATA ATG CGG TC |
| *Trim71(Exon2/3)* | TGA CAC CTG CTC TGT CCC CA | CAG TCT GGG CCT GCT CGA TG |
| *Lin28b* | CAT GGC ACT GGC CAC TGT AA | ATC ATG GAG ATG AAT CCG AAT CC |
| *Isl1* | CGG AGA GAC ATG ATG GTG GTT | AGG GCG GCT GGT AAC TTT G |
| *Rpl19* | GGT CTG GTT GGA TCC CAA | TGC CCG GGA ATG GAC AGT CA |

**Table S7 Antibodies and stains**

| **Reagent type** | **Designation** | **Source** | **Identifiers** | **Additional**  **information** |
| --- | --- | --- | --- | --- |
| antibody | TRIM71  rabbit polyclonal | gift from  Shinya Yamanaka | Gladstone Institute of Cardiovascular Disease | 1:500 dilution |
| antibody | MyosinVIIa  rabbit polyclonal | Proteus Biosciences | Cat.# 25-6790 | 1:500 dilution |
| antibody | SOX2  goat polyclonal | Santa Cruz | Cat.# sc-17320 | 1:500 dilution |
| antibody | Neurofilament  rabbit polyclonal | Millipore | Cat.# AB1989 | 1:1000 dilution |
| antibody | Parvalbumin  mouse monoclonal | Millipore | Cat.# P3088 | 1:500 dilution |
| antibody | CTBP2  Rabbit polyclonal | BioWorld Technology | Cat.#BS2287 | 1:500 dilution |
| antibody | β-actin  mouse monoclonal | Santa Cruz | Cat.# 47778 | 1:500 dilution |
| antibody | pSMAD2/3  rabbit polyclonal | Cell Signaling | Cat.# 8828 | 1:500 dilution |
| antibody | donkey anti-rabbit IgG (H+L) Alexa Fluor 647 | Thermo Fisher | Cat.# A-31573 | 1:1000 dilution |
| antibody | donkey anti-rabbit IgG (H+L) Alexa Fluor 555 | Thermo Fisher | Cat.# A-31572 | 1:1000 dilution |
| antibody | donkey anti-mouse IgG (H+L) Alexa Fluor 488 | Thermo Fisher | Cat.# A-21202 | 1:1000 dilution |
| antibody | Biotinylated donkey anti goat | Jackson Immuno Research Lab | Cat.#705-065-147 | 1:200 dilution |
| dye | Streptavidin, Alexa Fluor 405 | Life Technologies | Cat.#S32351 | 1:200 dilution |
| dye | Alexa Fluor 546 phalloidin | Invitrogen | Cat.#A22283 | 1:500 dilution |
| dye | Alexa Fluor 488 phalloidin | Invitrogen | Cat.#A12379 | 1:500 dilution |
| Nuclear stain | Hoechst 33258 solution | Sigma-Aldrich | Cat.# 94403 | 1:3000 dilution |
